# Chloroplast cpHsc70-1 interacts with VIPP1 C-terminal tail and controls VIPP1 oligomer disassembly in thylakoid membrane remodeling

**DOI:** 10.1101/2025.05.20.655051

**Authors:** Di Li, Sarah Wanjiru Gachie, Shin-Ichiro Ozawa, Martin Scholz, Michael Hippler, Wataru Sakamoto

## Abstract

Oxygenic photosynthetic organisms depend on the thylakoid membranes (TMs) for light-driven energy conversion. Recent studies on TM homeostasis (*thylakostasis*) have highlighted the essential role of the TM remodeling protein VIPP1 (Vesicle Inducing Protein in Plastids 1). As a member of the ESCRT-III/PspA/VIPP1 superfamily, VIPP1 forms large ring- and filament-like homo-oligomeric structures that exhibit a membrane remodeling activity. The oligomerization status was proposed to be modulated by the intrinsically disordered C-terminal tail (Vc), whereas its functional role remained unclear. Notably, this Vc region is conserved not only in photosynthetic VIPP1 but also in the PspA proteins of extremophilic species, implicating its role in membrane stress responses. To investigate the role of the Vc region in VIPP1 assembly, we performed co-immunoprecipitation assays in *Arabidopsis* chloroplasts and identified chloroplast-localized HSP70 proteins as major interactors. Among the two isoforms, cpHsc70-1 was found to be specifically required for modulating VIPP1 oligomeric assembly and dynamics in response to heat stress. Genetic analyses revealed that cpHsc70-1 facilitates the disassembly of VIPP1 oligomers, similarly to Vps4 ATPase in ESCRT-III; loss of either the Vc region or cpHsc70-1 impaired VIPP1 disassembly, resulting in more static oligomeric structures. Furthermore, cpHsc70-1 exhibited a broader role in chloroplast proteostasis, as the *cphsc70-1* mutant showed impaired accumulation of GFP-fusion proteins. Together, our findings uncover a crucial crosstalk between proteostasis and thylakostasis in chloroplasts, coordinated by cpHsc70-1 and VIPP1 in response to membrane stress.

**Significance Statement:** Organisms performing oxygen-evolving photosynthesis rely on the thylakoid membranes (TMs) for light-driven energy conversion. We reveal that VIPP1, a membrane remodeling protein forming dynamic oligomeric structures, is crucial for TM maintenance. Its intrinsically disordered C-terminal region (Vc) and the heat shock protein cpHsc70-1 coordinate VIPP1 disassembly during stress in *Arabidopsis* chloroplasts. Loss of either component causes defects in membrane remodeling and proteostasis. Our findings uncover a fundamental mechanism connecting membrane integrity and protein quality control, ensuring chloroplast resilience against environmental stress.

## Introduction

Oxygen-evolving photosynthetic organisms on Earth share a conserved mechanism for the light reaction, converting light energy into chemical energy (1). This reaction occurs exclusively within a specialized membrane called the thylakoid membrane (TM) (2). In the chloroplasts of vascular plants, the TM exhibits a complex architecture composed of stacked grana thylakoids and unstacked stroma thylakoids (3, 4). This spatial organization facilitates the distribution of the two photosystems, PSI and PSII, thereby promoting dynamic adaptation to fluctuating light environments (5, 6). The maintenance of TM homeostasis (*thylakostasis*) depends on the coordinated synthesis of lipid components (7, 8) and membrane shaping (9-13), the proper assembly of protein complexes (14, 15), and the regulation of protein complex assembly with post-translational modifications (3). In addition, membrane remodeling proteins have garnered attention as key regulatory factors in thylakostasis. Among them, Vesicle-Inducing Protein in Plastid 1 (VIPP1), a highly conserved member of the PspA/ESCRT-III/VIPP1 superfamily, has emerged as a central player.

VIPP1 is thought to have evolved from the bacterial Phage Shock Protein A (PspA) (16), which maintains membrane integrity in *E. coli* by preventing ion leakage (17). Both VIPP1 and PspA share conserved α-helical domains (H1–H6) (18-20), while VIPP1 possesses a unique C-terminal tail, H7, characterized as an intrinsically disordered region (IDR) (21). Originally identified as IM30 in pea(22), VIPP1 localizes to both the chloroplast inner envelope and the TM as large homo-oligomers (>2 MDa) in Arabidopsis (23, 24). VIPP1 has been implicated in a broad range of functions, including vesicle formation (13), membrane protection under stress (12), PSI supercomplex assembly in *Synechococcus* (25), PSII assembly during high-light stress in *Chlamydomonas* (26), and protein translocation across the TM (27). It binds preferentially to negatively charged lipids (28, 29) and promotes liposome aggregation and membrane fusion (27, 28, 30). Moreover, VIPP1 exhibits both ATPase and GTPase activity (31, 32).

While these findings highlight VIPP1’s multifunctionality, recent structural studies have emphasized its decisive role in membrane remodeling (33, 34). In its monomeric folding, H1 forms an amphipathic helix oriented vertically to the H2–H3 hairpin, while the C-terminal H4–H6 domains add flexibility and allow interweaving into symmetric oligomeric rings (13–18 protomers) that further assemble into layered structures reminiscent of a fruit basket. The disordered H7 tail faces the basket’s outer surface and remains structurally unresolved (35). Notably, VIPP1 shares structural similarity with not only PspA but ESCRT-III, a cytosolic component of ESCRT involved in endomembrane remodeling (35-37). In vitro AFM analyses revealed that VIPP1 from both cyanobacteria and *Chlamydomonas* self-assembles into diverse ESCRT-III-like structures—rings, carpets, and rods—on lipid bilayers without any additional factors (38, 39). The flexibility of VIPP1 monomers determines the relative abundance of these morphologies (39), which are thought to serve distinct roles in membrane remodeling and repair (40). In tobacco chloroplasts, VIPP1-GFP fusion proteins formed bundled filaments that underwent dynamic disassembly and reassembly in response to heat stress (41). These structures, termed functional VIPP1 particles (FVPs), are proposed to play key roles in thylakostasis.

To further understand the regulation of VIPP1 dynamics, we focused on the C-terminal H7 tail, hypothesizing that its disordered nature facilitates interactions with other proteins. Our previous in vivo studies using GFP-tagged VIPP1 constructs showed that deletion of the 43-amino-acid C-terminal region (Vc) led to larger FVP assembly and elevated sensitivity to heat stress, indicating that Vc negatively regulates VIPP1 self-assembly (21). In vitro AFM observation of cyanobacterial VIPP1 also demonstrated accelerated assembly upon the deletion of VIPP1 C-terminal regions, suggesting its involvement in oligomerization status (38). Interestingly, a recent *in silico* analysis revealed that not only photosynthetic VIPP1 proteins but also PspA proteins from extremophilic bacteria, such as Halobacteriales, possess Vc-like regions (Ma et al., manuscript under revision, included as Supplemental data). Although their sequences vary in length and composition, all Vc regions were predicted to be IDR, suggesting a conserved stress-responsive function, likely through protein–protein interactions.

Several approaches have been taken to characterize proteins interacting with VIPP1. In *Chlamydomonas*, it has been reported that the HSP70 chaperone comprised by HSP70B, cochaperone CDJ2, and nucleotide exchange factor CGE1 (42, 43), acts in both assembly and disassembly of VIPP1 rods in vitro (44-46). Detailed analysis of VIPP1/VIPP2 interactors through a proximity labeling method using biotin-based TurboID verified the interaction of the HSP70 chaperone with VIPP1 (47). In addition, a series of novel proteins (termed VIPP1 Proximity Labeling: VPL) potentially interacting with VIPP1/VIPP2 have been identified. These results demonstrate numerous proteins can interact with VIPP1, although the true function remains to be characterized. Interestingly, VPLs were revealed to be upregulated under stress conditions, implicating their roles in VIPP1 functionality and TM membrane remodeling. On the other hand, a recent work in Arabidopsis employed in affinity purification identified eight proteins interacting with VIPP1 (48). However, the possible relation of VIPP1 interactors to Vc is not well understood.

To further decipher VIPP1 interacting proteins and their function in vivo, we focused on Arabidopsis *vipp1* mutant complemented by expressing VIPP1-GFP. Co-immunoprecipitation and a series of genetic experiments demonstrated that chloroplast HSP70 (cpHsc70-1) is the main interactor through Vc, which controls VIPP1 oligomer disassembly rather than assembly. Our data thus demonstrates that in the chloroplast system, HSP70 is a disassembly factor of ESCRT-III oligomers substituting VPS4 ATPase in the cytosol.

## Results

### Search for proteins interacting with VIPP1 by GFP trap

To investigate proteins that interact with VIPP1, we employed transgenic lines expressing VIPP1 tagged by GFP at its C-terminus (Fig. 1*A*). We previously reported that either one of the transgenes, expressing VIPP1-GFP or VIPP1ΔC-GFP) lacking Vc, rescues the lethality of null *vipp1* mutant (*vipp1-ko*), indicating that VIPP1-GFP is functional and that Vc is dispensable under normal growth conditions (21). Chloroplast proteins purified from these two lines, *VIPP1-GFP/vipp1-ko* (*VIPP1-GFP/ko*) and *VIPP1ΔC-GFP/vipp1-ko* (*VIPP1ΔC-GFP/ko*) were used in this study to perform GFP trap coimmunoprecipitation (TurboGFP-Trap® Magnetic Agarose). Structurally, Vc faces the outer surface of VIPP1 rings, but its own structure was not resolved due to its intrinsically disordered property (21, 35). In situ observation of GFP signals showed that VIPP1ΔC-GFP formed highly interconnected VIPP1 spheres and rods (Fig. 1*B*), demonstrating its role in negatively regulating VIPP1 oligomer formation and dynamics responding to osmotic stress (21). It is conceivable that some of the chloroplast proteins interact with VIPP1 via Vc.

**Fig. 1.**
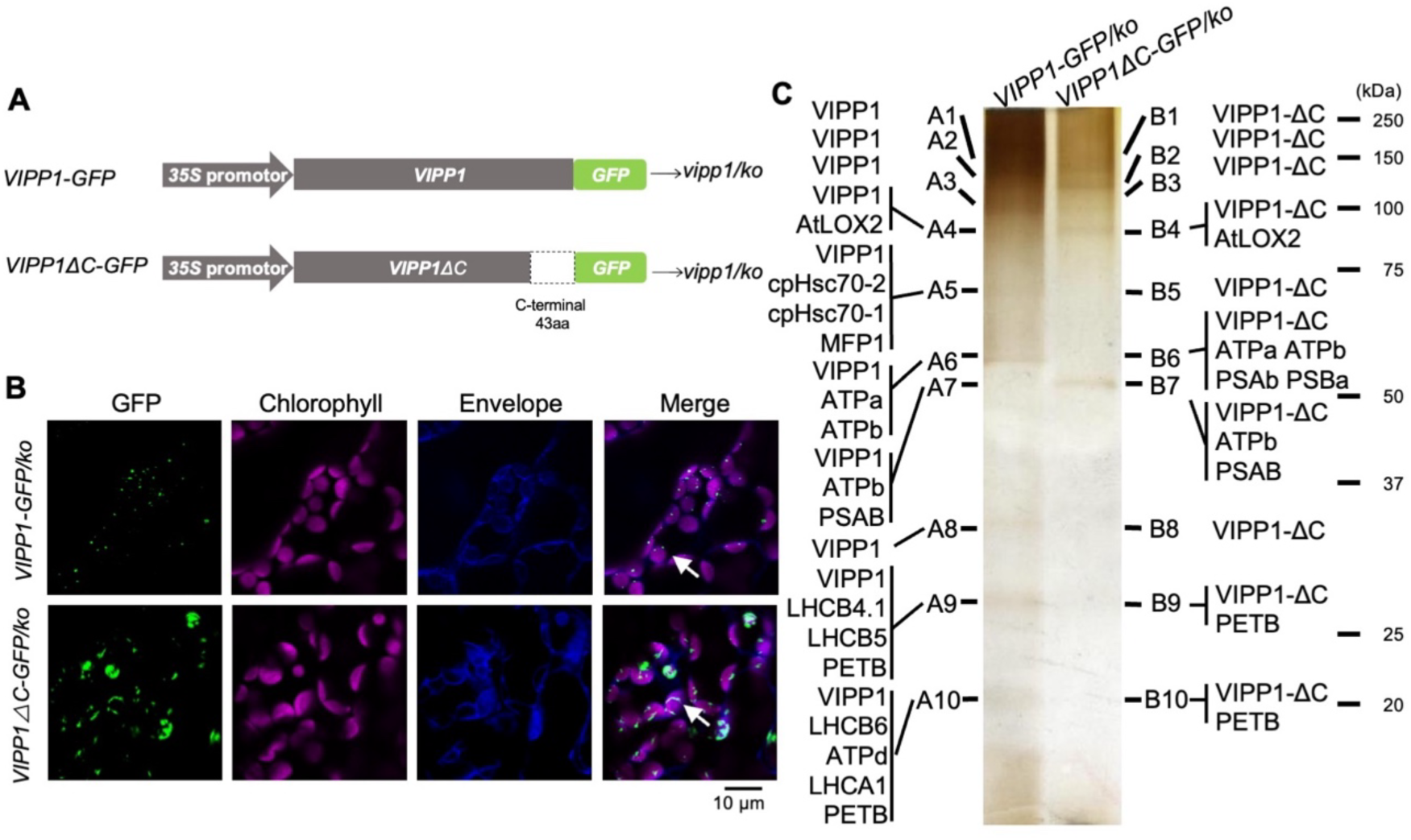
Arabidopsis *VIPP1* transgenic lines used in this study and immunoprecipitation analysis using GFP trap. (A) Schematic representation of the *V/PP1-GFP* and *VIPP1ΔC-GFP* transgenes introduced into *vipp1-ko.* GFP (green) was fused to the C-terminus of VIPP1 or VIPP1*L’>C* lacking Ve (grey). Expression both constructs was driven by the CaMV *35S* promoter (arrows). Deletion of Ve in *VIPP1ΔC-GFP* is indicated by an open box. (B) Representative confocal images of *VIPP1-GFP/ko* and *VIPP1ΔC-GFP/ko* in true leaves. GFP fluorescence, chlorophyll autofluorescence, and envelope stained by Rhodamine B are shown individually and as merged images. Bar= 10 µm. (C) SOS-PAGE of proteins immunoprecipitated by GFP trap. Solublized chloroplast protein extracts from *VIPP1-GFP* and *VIPP1ΔC-GFP* were immunoprecipitated using anti-GFP serum-coupled beads. Samples were separated by SOS-PAGE and visualized by silver staining. Gel slices from *VIPP1-GFP* (A1-A10) and *VIPP1ΔC-GFP* (B1-B10) were excised from the gels and analyzed by mass spectrometry. Identified proteins are listed in Supplemental Dataset 1.

Chloroplast proteins solubilized by 1% Dodecyl-β-D-Maltopyranoside (DDM) were incubated with GFP-trap agarose beads, and the bound fraction was resolved by SDS-PAGE (Fig. 1*C*). Silver stain of the gels detected several bands (A1 to A10 in *VIPP1-GFP/ko* and B1 to B10 in *VIPP1ΔC-GFP/ko*), including those that corresponded to VIPP1-GFP (A6) and VIPP1ΔC-GFP (B7). All detected bands were excised from the gel and subjected to (MS) (Suppl Dataset 1). As expected, VIPP1 peptides were the most abundant in A6 and B7. VIPP1 was also detected in all the excised bands, likely due to its abundance in the immunoprecipitate. The proteins with the highest abundance in each band are annotated (Fig. 1*C*, a full list of identified proteins is presented in Suppl Dataset 1).

### Chloroplast HSP70 and MFP1 as candidates interacted with VIPP1

While most of the bands included proteins that correspond to structural components of photosystems, light-harvesting antennae and ATPase in the TM, A5 was specific to *VIPP1-GFP/ko* and contained three unique proteins (cpHsc70-1, cpHsc70-2, MFP1). B5 is the corresponding region in *VIPP1ΔC-GFP/ko* and had only VIPP1ΔC peptides. cpHsc70-1 and cpHsc70-2 are the two HSP70 isoforms in chloroplasts (cpHSP70) (49-51). MFP1, well conserved in a wide range of dicotyledonous and monocotyledonous plants, was reported to be localized in TMs (52). Recently, MFP1 was shown to affect the location and quantity of starch formation in chloroplasts (53). It is noteworthy that these two proteins were missing in B5 and any other bands from *VIPP1ΔC-GFP/ko*. In our triplicated experiments, cpHSP70 and MFP1 were reproducibly detected in *VIPP1-GFP/ko* (A1 in Fig. S1 and A2 in Fig. S2; Suppl dataset 2 and 3). Thus, we considered that cpHSP70 and MFP1 were the candidates to interact with VIPP1 through Vc. In plants, HSP70 is regulated by biotic stress (54, 55) and known to assist in various cellular processes, including protein quality control, transmembrane transport, protein degradation, and intracellular signal transduction (51, 56, 57). In *Chlamydomonas*, Hsp70B was shown to interact with VIPP1 (44, 45). Given these functions related to VIPP1 oligomerization dynamics, we focused on cpHsc70-1 and cpHsc70-2 further in this study.

### Verification of interaction between VIPP1 and cpHsc70 by co-immunoprecipitation

To verify the interaction between cpHsc70 and VIPP1, co-immunoprecipitation was performed. We first confirmed our GFP trap assay using anti-GFP-coupled Magnosphere beads. The results were then visualized by Western blots immunoreacted with anti-VIPP1 and anti-cpHSP70 antibodies (derived from pea stromal HSP70 S78, recognizing cpHsc70-1 and cpHsc70-2). As expected, the VIPP1 antibody recognized a single band in both *VIPP1-GFP/ko* and *VIPP1ΔC-GFP/ko* eluates (Fig. 2*A*). Consistent with our MS data, cpHSP70 was detected as a single band in *VIPP1-GFP/ko* but not *VIPP1ΔC-GFP/ko* eluates (Fig. 2*A* and *E*). We also performed coimmunoprecipitation using Magnosphere beads conjugated with the cpHSP70 antibody. While chloroplast proteins immunoprecipitated cpHSP70 successfully, VIPP1 was co-immunoprecipitated only in *VIPP1-GFP/ko* but not in *VIPP1ΔC-GFP/ko* (Fig. 2*B* and *F*).

**Fig. 2.**
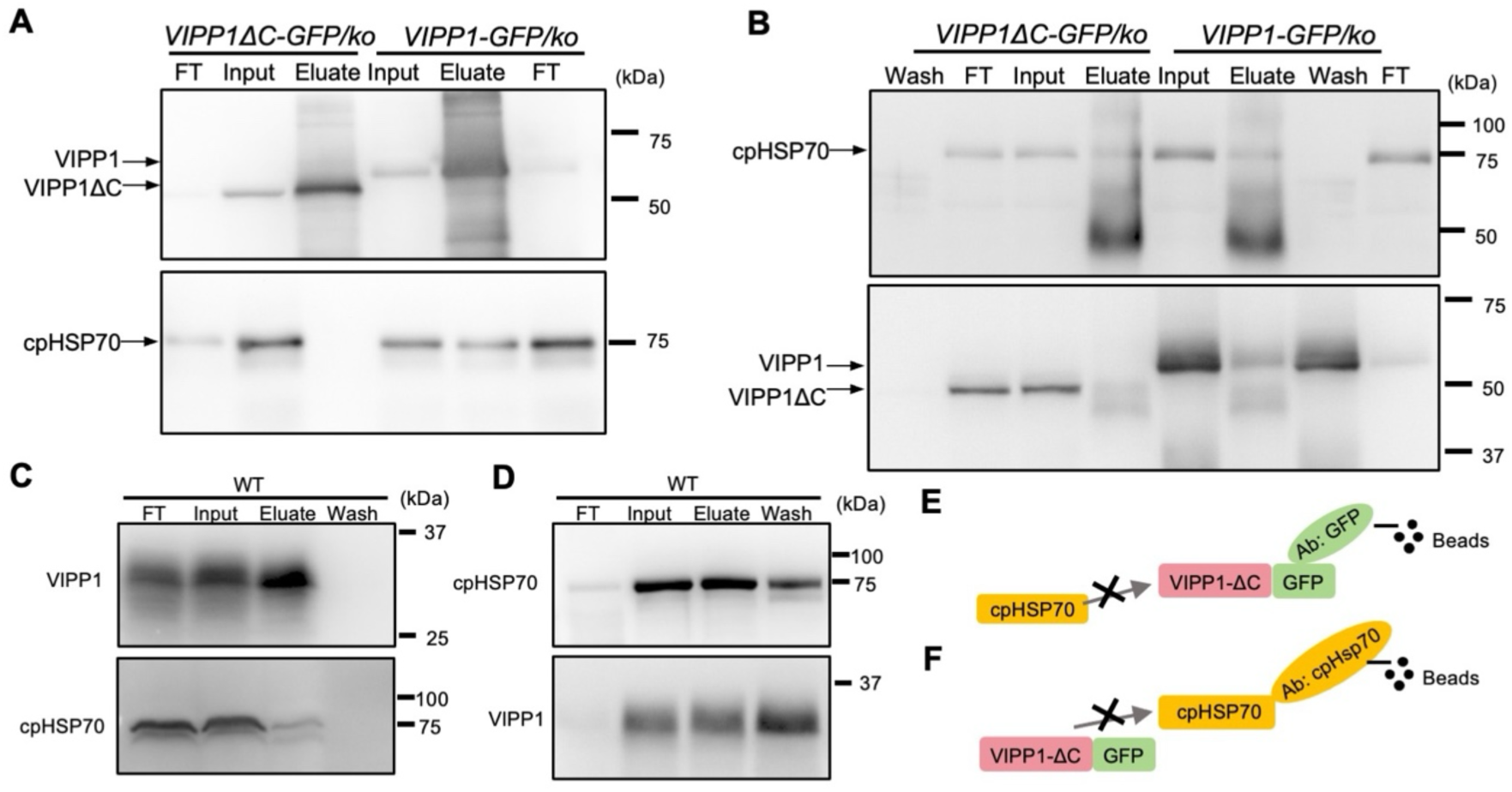
Immunoprecipitation of cpHSP70 with VIPP1-GFP and endogenous VIPP1. (A) Validation of GFP-trap results. Soluble extracts from *VIPP1-GFP/ko* and *VIPP1ΔC-GFP/ko* were incubated with anti-GFP-coupled beads. lmmunoprecipitated proteins were analyzed by immunoblotting using anti-VIPP1 (top) and anti-cpHSP?O (bottom) antibodies. (B) Co-immunoprecipitation with anti-cpHSP?O antibodies. As in (A), but immunoprecipitation was carried out using anti-cpHsp?O-coupled beads. Eluted proteins were probed with anti-cpHsp?O (top) and anti-VIPP1 (bottom) antibodies. (C, D) Co-immunoprecipitation in WT plants. Total proteins extracts from WT were incubated with anti-VIPP1-coupled beads (C) or anti-cpHsp?O-coupled beads (D). Precipitated proteins were probed with anti-VIPP1 and cpHsp?O antibodies, as indicated. (E, F) Schematic illustration of co-immunoprecipitation results using anti-GFP-coupled beads (E) and anti-cpHSP70-coupled beads. No interaction was observed between VIPP1ΔC-GFP and cpHSP70 in either experiment (see (A) and (B)).

Co-immunoprecipitation was also conducted in the wild-type Columbia (Col) plant. Magnosphere beads conjugated with either VIPP1 or cpHSP70 antibody were prepared for the co-immunoprecipitation assays. Immunoblot analysis revealed that when using VIPP1 antibody-conjugated beads, cpHSP70 was co-precipitated by VIPP1 *in vivo* (Fig. 2*C*). Reciprocally, when using cpHSP70 antibody-conjugated Magnosphere beads, VIPP1 was also pulled down *in vivo* (Fig. 2*D*).

### Verification of interaction between VIPP1 and cpHsc70 by His tag

To ascertain that the interaction between cpHSP70 and VIPP1 is not influenced by the presence of the GFP tag, we created another transgenic line in which *VIPP1* was C-terminally tagged by the sequence corresponding to 6xHis (Fig. 3*A*) and introduced it into *vipp1-ko* mutant background. The resulting transgenic line *VIPP1-His/ko* grew comparably to Col and exhibited no detectable phenotypes (Fig. 3*B*), demonstrating full complementation of *vipp1-ko*. Similarly to the wild-type VIPP1 and VIPP1-GFP, VIPP1-His forms large oligomers whose size estimated by Blue-Native PAGE was greater than photosynthetic complexes in the TMs (Fig. 3*C*). We next purified VIPP1-His using Ni-NTA column (Fig. 3*D*). The purified fraction was subjected to mass spectrometry (Suppl dataset 4), in which 379 proteins were detected as those found in *VIPP1-His/ko* but not in the negative control derived from Col (Suppl Dataset 4). Importantly, both cpHsc70-1 and MFP1 were included in this dataset. Our western blot indeed confirmed that VIPP1-His purified from Ni-NTA column also contained cpHSP70 (Fig. 3*E*). Taken together, these results led us to conclude that cpHSP70 interacts with VIPP1. As presumed from the structural analysis, C-terminally tag to VIPP1 appeared to have no negative effect on protein interactions, whereas no detectable signal of cpHSP70 in *VIPP1ΔC-GFP/ko* eluates suggested that Vc is required for the interaction between VIPP1 and cpHSP70.

**Fig. 3.**
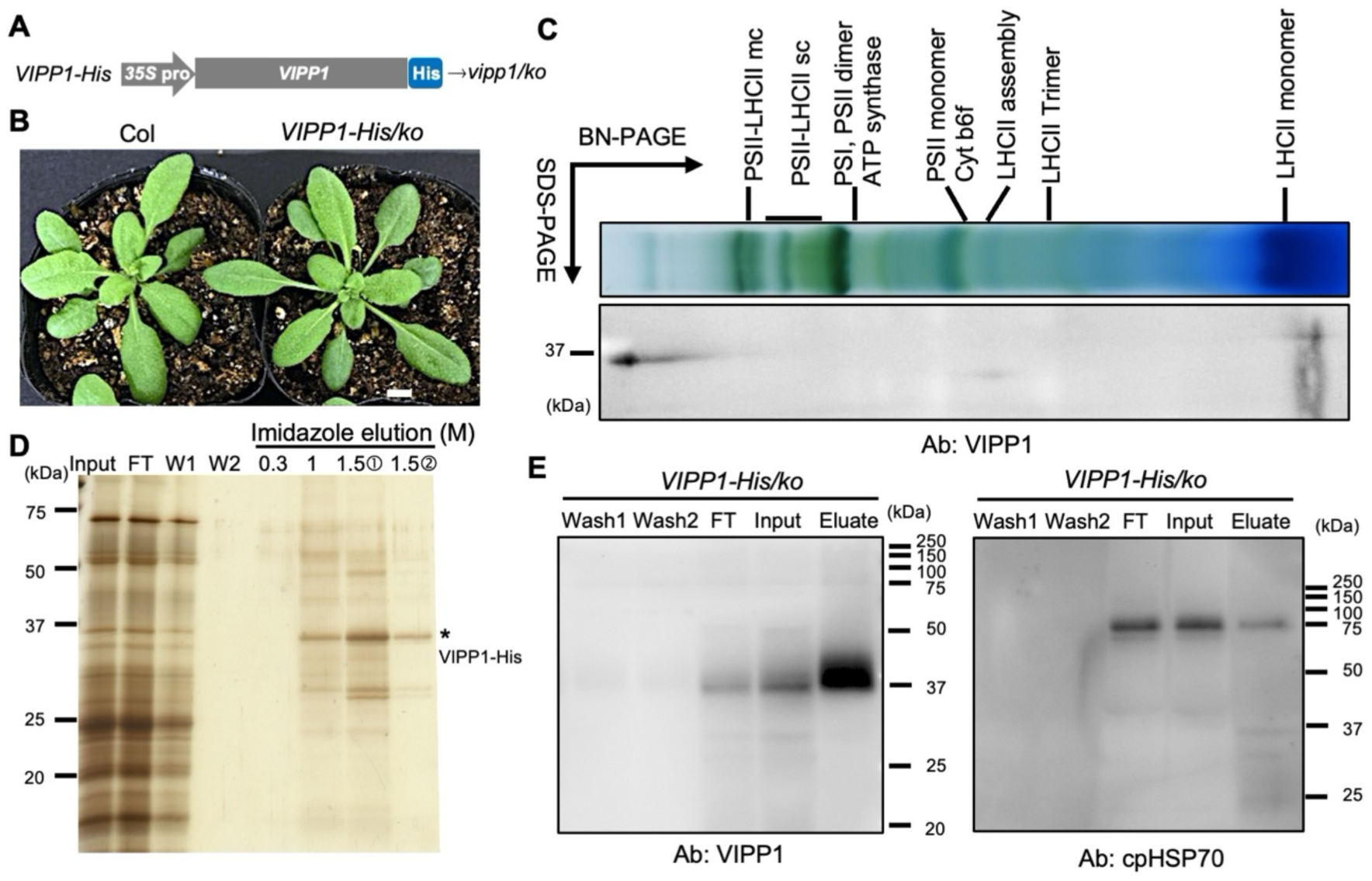
In vivo interaction between VIPP1 and cpHSP70 confirmed in transgenic plants expressing VIPP1-His. (A) Schematic representation of the *VIPP1-His* transgene introduced into *vipp1-ko,* driven by CaMV *35S* promoter *(35S* pro). (B) Morphology of wild-type (Col) and a representative *VIPP1-Hislko* line, showing full phenotypic complementation of the *vipp1-ko* mutation. Bar= 1 cm. (C) Buie-native (BN) PAGE (first dimension) followed by SOS-PAGE (second dimension) analysis of thylakoid membrane proteins solubilized by 1% DOM. The upper panel shows a BN-PAGE gel slice displaying the separation of photosynthetic complexes (labeled at the top). The lower panel shows a western blot of the second-dimension gel probed with VIPP1 antibodies. (D) Purification of VIPP1-His using Ni-NTA column. Input: total input of thylakoid proteins, FT: flow through fraction, W1 and W2: washed fractions. Eluted fractions with increasing imidazole concentrations (0.3, 1, and 1.5 **M** first and second) are shown. The VIPP1-His band is marked with an asterisk. Proteins identified by mass spectrometry are listed in Supplemental Dataset 4. (E) lmmunoblots of the 1 M imidazole-eluted fractions from (D), probed with anti-VIPP1 and anti­ cpHSP70 antibodies.

### Thermosensitivity of *cphsc70-1* seedlings was phenocopied in *VIPP1ΔC-GFP/ko* but not *VIPP1-GFP/ko*

Mutants lacking *cpHsc70-1* and *cpHsc70-2* have been characterized previously, where *cphsc70-1* displayed weak growth at the seedling stage. Both *cphsc70-1* and *cphsc70-2* were viable and set seeds, whereas the double mutant is embryo-lethal (49). Although two proteins are highly homologous and accumulate substantially in vegetative growth stage, cpHsc70-1 appears to be the major isoform in chloroplasts. Consistent with this supposition, *cphsc70-1* exhibited elevated thermosensitivity against heat shock, compared to the wild type and *cphsc70-2* plants under our growth conditions(49, 58).

To investigate the effects of impaired interaction between VIPP1 and HSP70, heat treatment experiments were conducted on the seedlings from *VIPP1-GFP/ko, VIPP1ΔC-GFP/ko, cphsc70-1*, and *cphsc70-2*, along with the control Col and *VIPP1-GFP/Col* plants. We followed the heat-shock condition engaged essentially in the previous work as summarized in Fig. 4*A*. After pre-heat treatment and subsequent acclimation for 2 days, seedlings were exposed to heat shock and incubated further at growth temperature for 5 days (Fig. 4*B*). While we reproduced paler phenotypes of *cphsc70-1* seedlings as previously reported, *VIPP1ΔC-GFP/ko* displayed noticeably paler phenotype similar to *cphsc70-1*, as revealed by chlorophyll measurement (Fig. 4*C*). These results indicated that the thermosensitivity caused by the lack of cpHsc70-1 could be phenocopied in *VIPP1ΔC-GFP/ko* but not *VIPP1-GFP/ko*, strongly suggesting the interaction between cpHsc70-1 and Vc against heat stress. To examine if the observed thermosensitivity is correlated with cpHsc70 levels, we performed western blotting under the heat stress. However, no significant increase of cpHSP70 was detected in Col (Fig. 4*D*), although assessments of transcript levels in the public database indicated that *cpHsc70-1* and *cpHsc70-2* transcripts increased after heat shock (Fig. S3). The protective role of VIPP1 under heat stress does not result from mere upregulation of cpHsc70 and VIPP1.

**Fig. 4.**
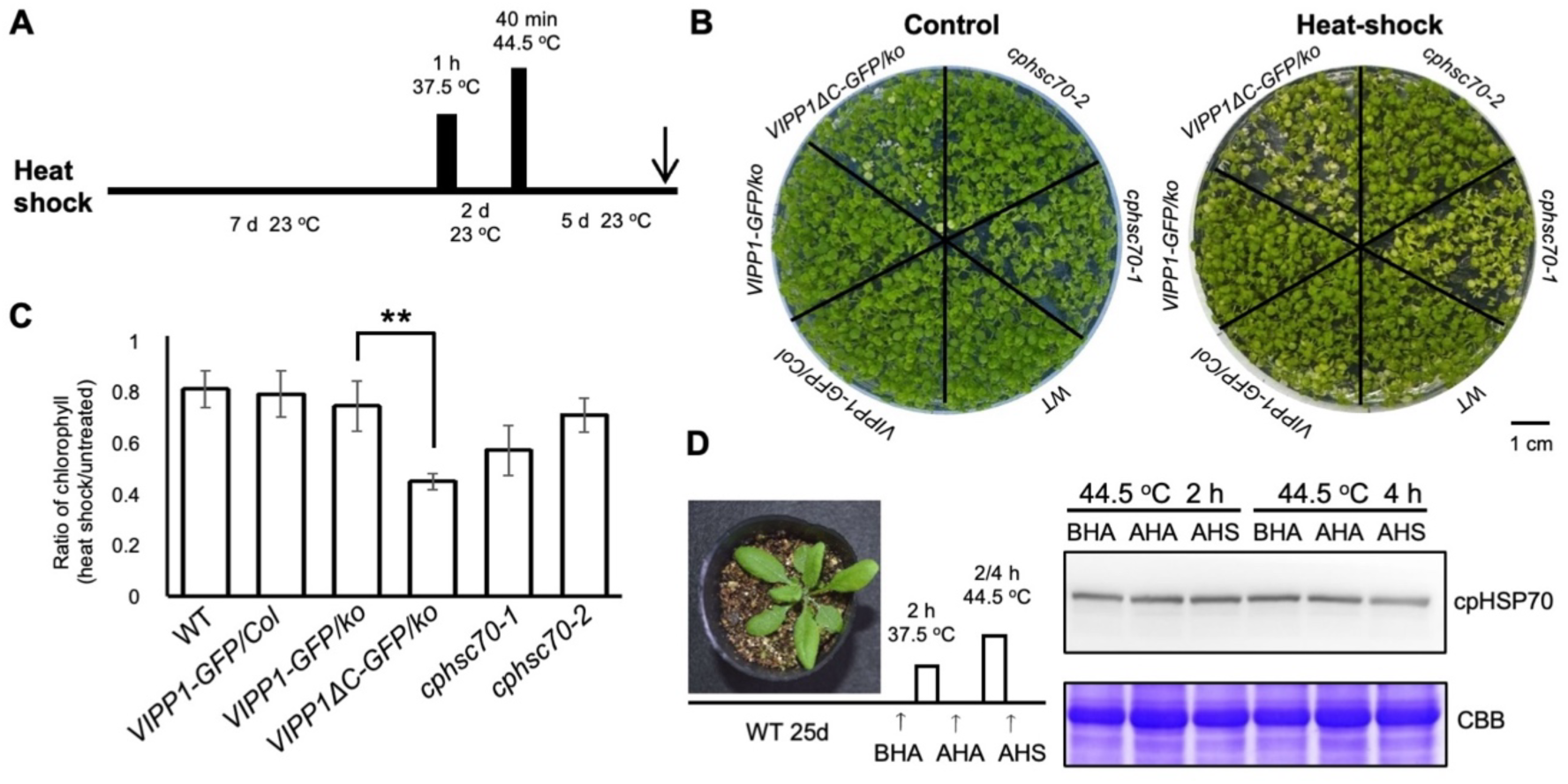
Assessment of heat shock tolerance in cpHSP70 mutants and VIPP1 transgenic lines. (A) Schematic diagram of experimental design. Seven-day-old seedlings were subjected to a heat acclimation treatment at 37.5 °C for 1 hour, followed by a 2-days recovery period at 23 °C. Plants were then exposed to heat shock at 44.5 °C for 40 minutes, and the phenotypes were assessed after a 5-day recovery period (indicated by the arrow). Control plants were continuously grown at 23 °C. (8) Phenotypes of control (left) and heat-treated (right) seedlings of WT, *cphscl0-1, cphscl0-2, VIPP1-GFP/ko, VIPP11ΔC-GFP/ko,* and *VIPP1-GFP/Co/.* A representative image from three biological replicates is shown. (C) Ratios of chlorophyll content in heat-treated plants relative to untreated controls (N=3). Asterisks denote significant difference at p<0.01 (Student’s t-test). (D) cpHsp70 protein levels in WT during heat treatment. Twenty-five-day-old WT plants were subjected to the heat shock regimen shown in (A), and total protein was extracted from three time points: before heat acclimation (BHA), after heat acclimation (AHA), and after heat shock (AHS). A representative western blot (from two biological replicates) was probed with anti-cpHsp70 antibodies. Protein leading was normalized based on chlorophyll content and visualized with Coomassie Brilliant Blue (CBB) staining.

### Genetic study 1: cpHsc70-2 has no effects on VIPP1 oligomeric status and dynamics

To elucidate the effects of cpHsc70 on VIPP1 oligomeric status and dynamics, we attempted to make a cross to generate *VIPP1-GFP/ko* and *VIPP1ΔC-GFP/ko* under *cphsc70-1* or *cphsc70-2* backgrounds. While we faced difficulty to create *VIPP1-GFP* or *VIPP1ΔC-GFP* strains under the *vipp1-ko* and *cphsc70-1* double-mutant backgrounds (see below), we successfully obtained *VIPP1-GFP/vipp1-ko/cphsc70-2* (*cphcs70-2 VIPP1-GFP/ko*) and *VIPP1ΔC-GFP/vipp1-ko/cphsc70-2* (*cphsc70-2 VIPP1ΔC-GFP/ko*) lines (Fig. S4 *A* and *B*). These two lines grew normally and showed no visible phenotypes (Fig. S5*A*). GFP fluorescence in these lines was compared with the original *VIPP1-GFP/ko* and *VIPP1ΔC-GFP/ko* lines, in which the number and length of GFP particles (termed FVPs: functional VIPP1 particles) were significantly different between the two lines (Fig. S5 *B, C* and *D*); namely, upon the removal of Vc, the robust assembly of FVPs became apparent (Fig S5*B*), and its dynamics are hindered when compared with VIPP1-GFP (Fig. S5*E*, Suppl Video *1* and *3*).

The majority of FVPs exhibited mobility and moved with different speed. In contrast, VIPP1ΔC-GFP exhibited large filaments and various structures that are virtually stationary. When *cpHsc70-2* was knocked out, this difference between VIPP1-GFP and VIPP1ΔC-GFP showed no significant change (Fig S5*B*). In *cphsc70-2 VIPP1-GFP/ko*, the morphology and dynamics of FVPs appeared to be indistinguishable from those in *VIPP1-GFP/ko* chloroplasts (Fig. S5*B*). Similarly, in *cphsc70-2 VIPP1ΔC-GFP/ko*, VIPP1ΔC-GFP exhibited larger and sometimes filamentous assemblies (Fig. S5*B*). VIPP1-GFP showed noticeable mobility whereas VIPP1ΔC-GFP remained comparatively inert and immobile (Fig. S5*B*, Suppl Video *5* and *6*). Together, these results indicated that cpHsc70-2 has no or little effects on VIPP1 oligomeric status and dynamics.

### Genetic study 2: proper expression of VIPP1-GFP requires cpHsc70-1

We originally attempted to generate *cphsc70-1 VIPP1-GFP/ko* and *cphsc70-1 VIPP1ΔC-GFP/ko* lines, similarly to the case for *cphsc70-2* as aforementioned. To do this, we crossed *cphsc70-1* to *VIPP1-GFP/ko* or *VIPP1ΔC-GFP/ko.* Despite multiple rounds of F2 screening, however, no such double mutants were obtained (Fig. 5*A, B, C,* and *D*); screening the progeny harboring *VIPP1-GFP* (or *VIPP1ΔC-GFP*) and exhibiting the leaf phenotype of *cphsc70-1* in all F2 and the following generations resulted in either *vipp1-ko* +/+ (wild type) or *vipp1-ko* +/-(heterozygous) genotype. No individuals homozygous for *vipp1-ko* -/- was obtained (Fig. 5 *A* and *B*, Fig. S4 *C* and *D*). To confirm this result, we selected an F2 individual showing the presence of *VIPP1-GFP* or *VIPP1ΔC-GFP* with *cphsc70-1* -/- and *vipp1* +/-genotypes, and the segregation of *vipp1* was investigated in the subsequent F3 population (Fig. 5*C* and *D*). Consequently, no plants showing *vipp1-ko* -/- were obtained, whereas *vipp1-ko* +/- and *vipp1-ko* +/+ genotypes segregated at the ratio of 2:1 (Fig 5 *C* and *D*). Observation of seed setting in the silique of the selfed plants indeed showed that a portion of seed settings was aborted, indicating the embryo lethality of the *vipp1-ko cphsc70-1* double mutant (Fig. 5 *E* and *F*). Together, these data demonstrated that the rescue of *vipp1-ko* lethality by expressing VIPP1-GFP or VIPP1ΔC-GFP requires cpHsc70-1.

**Fig. 5.**
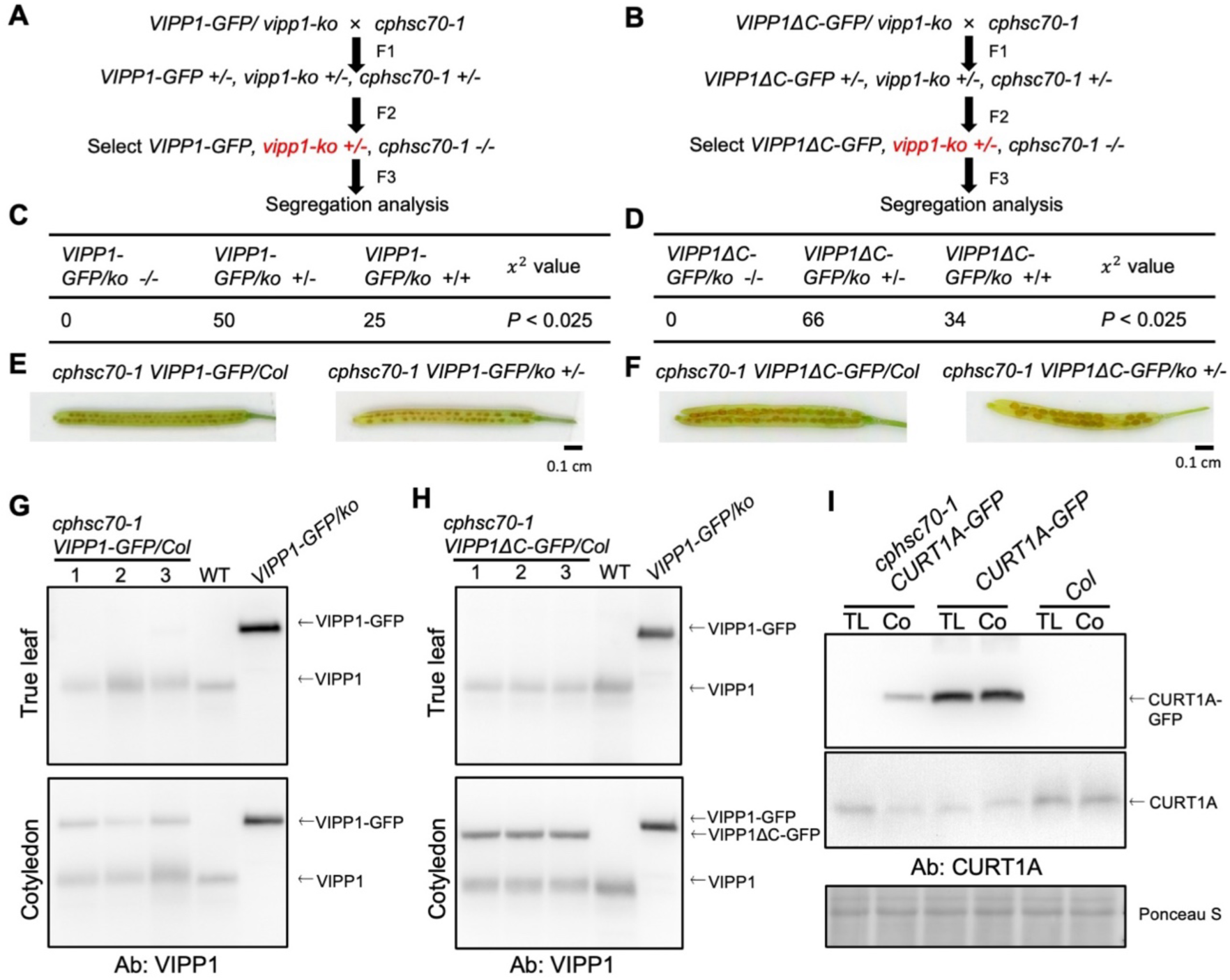
Segregation analysis of *VIPP1-GFP/ko* +/- and *VIPP1ΔC-GFP/ko* +/-genotypes in *cphscl0-1* background reveals the requirement of cpHsc70-1 for expression of GFP-tagged VIPP1 proteins. (A, 8) Diagrams showing the genetic analysis performed in *VIPP1-GFP/ko* x *cphscl0-1* (A) and *VIPP1ΔC-GFP/ko* x *cphscl0-1* (B) crosses. In each crossed population, an F2 individual homozygous for the GFP-tagged *VIPP1* allele and *cphscl0-1* but heterozygous for *vipp1* (indicated in red), was selected. Segregation of the *vipp1* +/-in the subsequent F3 generation was examined. (C, D) Segregation from the F3 populations in (A) and (8), showing the absence of *vipp1-ko* -/- progeny, suggesting a lethal interaction. (E, F) Representative siliques dissected from plants of the indicated genotypes. The results demonstrate that the coexistence of GFP-tagged VIPP1, *vipp1,* and *cphsc70-1* is lethal. (G) Accumulation of VIPP1 proteins in true leaves (top) and cotyledons (bottom) of *cphscl0-1 VIPP1-GFP/Co/* (three biological replicates: 1, 2, and 3), along with WT and *VIPP1-GFP/ko* as controls. Bands detected with anti-VIPP1 antibody are indicated with arrows. (H) Accumulation of VIPP1 proteins in true leaves (top) and cotyledons (bottom) of *cphscl0-1 VIPP1ΔC-GFP/Co/* (three biological replicates: 1, 2, and 3), along with WT and *VIPP1-GFP/ko* as controls. Bands detected with anti-VIPP1 antibody were indicated with arrows. (I) Accumulation of CURT1A proteins in true leaves (TL) and cotyledons (Co) of *cphscl0-1 CURT1A-GFP* (left), *CURT1A-GFP/Co/* (middle), and Col (right). Bands corresponding to CURT1A-GFP (top) and CURT1A (middle) are indicated with arrows. Protein loading was normalized to chlorophyll content, and Ponceau S-stained membrane is shown as a loading control (bottom).

To rationalize the lethality of *VIPP1-GFP/ko* or *VIPP1ΔC-GFP/ko* without cpHsc70-1, we raised the possibility that accumulation of both VIPP1-GFP and VIPP1ΔC-GFP was impaired in the *cphsc70-1* background. To examine this possibility, we performed western blotting using the VIPP1 antibody. While our Western blots barely detected VIPP1-GFP or VIPP1ΔC-GFP in true leaves of both *cphsc70-1 VIPP1-GFP/Col/* or *cphsc70-1 VIPP1ΔC-GFP/Col* (Fig. 5 *G* and *H*), the band corresponding to VIPP1-GFP or VIPP1ΔC-GFP was detected in cotyledons (Fig. 5 *G* and *H*). These results suggested that the stable accumulation of VIPP1-GFP proteins requires cpHsc70-1, regardless of the presence or absence of Vc.

To assess whether the effect of the *cphsc70-1* mutation is specific to VIPP1 or rather common to other proteins, we decided to investigate another TM remodeling protein CURT1A (9). We generated transgenic plants expressing CURT1A-GFP in the wild-type background (*CURT1A-GFP/Col*), in which steady accumulation of CURT1A-GFP was detected by Western blotting using anti-CURT1A antibody (Fig. 5 *I*). However, when we created this transgenic line under the *cphsc70-1* background (*cphsc70-1 CURT1A-GFP*), accumulation of CURT1A-GFP was compromised as observed in the case of VIPP1; CURT1A-GFP was detectable in cotyledons but not in true leaves (Fig. 5 *I*). These results suggested that cpHsc70-1 has profound effects on the proper accumulation of GFP-tagged proteins. This effect of cpHSC70-1 on the GFP was not confined to VIPP1 but rather appeared to be widespread in chloroplast proteins (59).

### Genetic study 3: cpHsc70-1 is required for the proper function of VIPP1 by preventing excess oligomeric assembly

To explore the effect of cpHsc70-1 in VIPP1 dynamics, we next focused on *cphsc70-1 VIPP1-GFP/Col* and *cphsc70-1 VIPP1ΔC-GFP/Col* further. These transgenic lines were viable and set seed normally with the exception that they exhibited a similar phenotype as *cphsc70-1* (49, 56), displaying curled leaf margin (Fig. 6 *A*, Fig. S4 *A*). We next observed GFP signals in these two lines. Consistent with Western blotting results, no GFP signals were observed in true leaves, whereas they were detectable in cotyledons, although less to the control *VIPP1-GFP/Col* or *VIPP1ΔC-GFP/Col* plants. Observation of GFP signals by confocal microscopy revealed that the knockout of cpHsc70-1 profoundly impacted the morphology of FVPs, presenting elongated rod-like GFP signals which were similar to those in *VIPP1ΔC-GFP/ko* (Fig. 6 *B* and *C*). Quantitative analysis of VIPP1-GFP signals revealed that the number of GFP particles per chloroplast decreased significantly in *cphsc70-1 VIPP1-GFP/Col* (Fig. 6*E*). The length of GFP particles also showed significant increase (Fig. 6*F*). In contrast, GFP signals in *cphsc70-1 VIPP1ΔC-GFP/Col*, however, did not exhibit any significant phenotypic alterations when compared to *VIPP1ΔC-GFP/ko* (Fig. 6 *B* and *C*). As for the dynamics of GFP particles, *cphsc70-1 VIPP1-GFP/Col* displayed retarded movement compared to *VIPP1-GFP/ko*, although the defective movements were not to the extent observed in *VIPP1ΔC-GFP/ko* (Fig. 6 *G*, Suppl Video *2*, *4* and *7*). In contrast, VIPP1ΔC-GFP remained relatively immobile irrespective of the presence or absence of cpHsc70-1 (Fig 6 *G*, Suppl Video *4* and *8*).

**Fig. 6.**
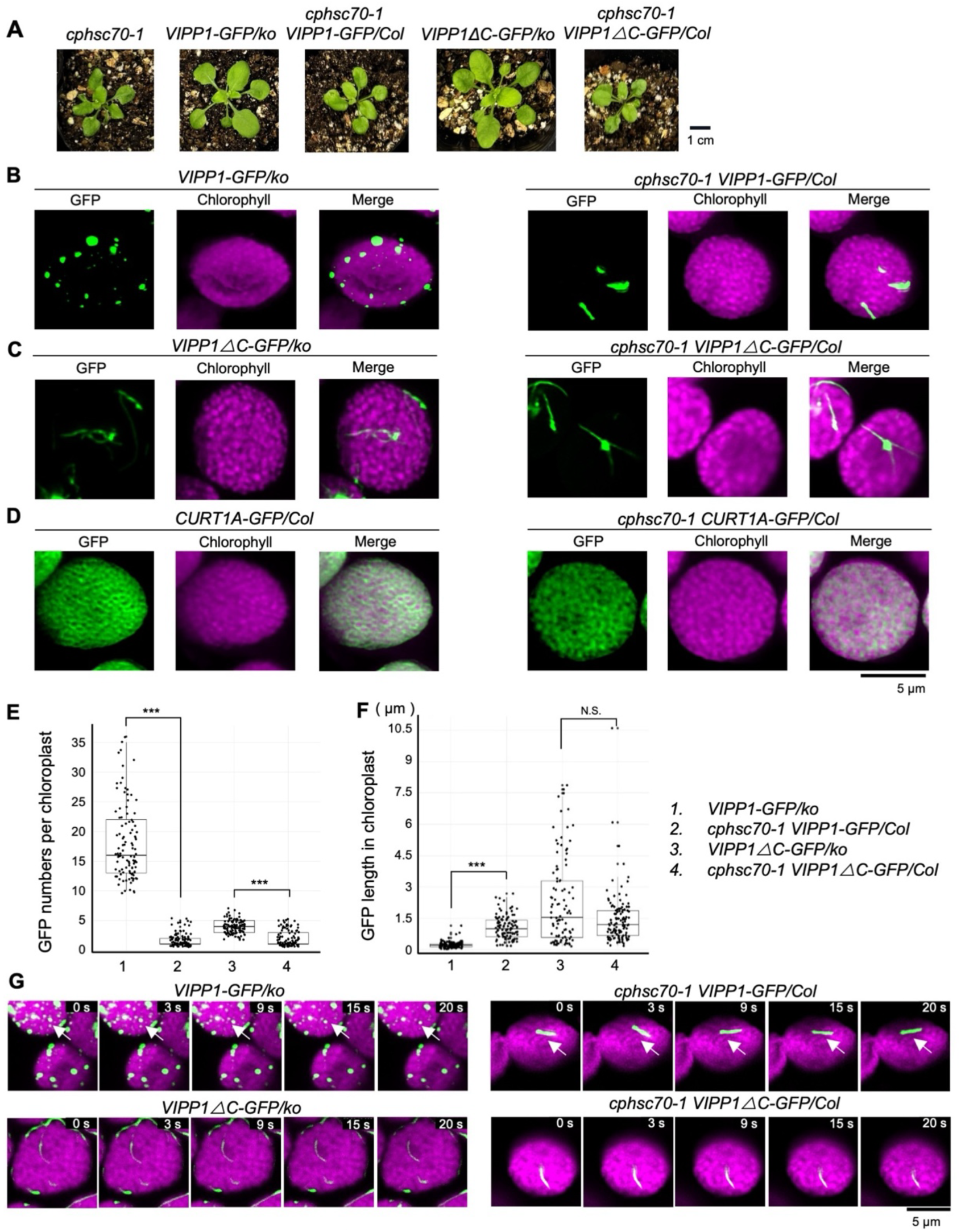
Confocal microscopic observation of GFP-tagged VIPP1 proteins in chloroplasts. (A) Plant morphology at 15 days after germination. Genotypes are indicated above the images. (B) A representative image of VIPP1-GFP in cotyledon chloroplasts from *vipp1-ko* (left) or *cphsc70-1* (right) background. (C) A representative image of VIPP1ΔC-GFP in cotyledon chloroplasts from *vipp1-ko* (left) or *cphsc70-1* (right) background. (D) A representative image of CURT1A-GFP in cotyledon chloroplasts from Col (left) or *cphsc70-1* (right) background. GFP, chlorophyll autofluorescence, and merged images are shown for each. Bar=5 µm. (E) Box plots showing the number of GFP particles per chloroplast for each genotype (N > 105). (F) Box plots showing the length (longest axis) of indivicual GFP particles, analyzed by image J (N > 105). (G) Time lapse images of GFP-tagged VIPP1 particles in cotyledons from *VIPP1-GFP/ko* (top left), *VIPP1ΔC-GFP/ko* (bottom left), *cphsc70-1 VIPP1-GFP/Co/* (top right), *and cphsc70-1 VIPP1L’.C­ GFP/Co/* (bottom right). Shown are five time points (0-20 seconds) from each time-lapse series. White arrows indicate fast moving GFP particles.

To examine if the effect of the *cphsc70-1* mutation on the assembly or spatial organization of GFP-tagged proteins is seen in other proteins than VIPP1, we observed GFP signals in *CURT1A-GFP* under either Col or *cphsc70-1* background. As involved in membrane curvature of TMs, CURT1A has been shown to be localized in curvature domains. Consistent with this, GFP signals were observed in the region surrounding grana discs, represented by chlorophyll autofluorescence (Fig. 6 *D*). The spatial organization of CURT1A-GFP was comparable in Col and *cphsc70-1* background. Together with a series of genetic experiments, our data demonstrated that cpHsc70-1 negatively regulates the oligomerization status of FVPs, as the lack of cpHsc70-1 resulted in forming larger filamentous FVPs. Given that similar filamentous FVPs were observed in *VIPP1-GFP/ko* under *cphsc70-1* background or in *VIPP1ΔC-GFP* lacking Vc, the dynamic behavior of VIPP1 oligomers responding to heat and other stresses appeared to be regulated by the interaction of Hsc70-1 with VIPP1 at Vc.

## Discussion

### Identification of proteins interacting with the Vc region

In this study, we focused on the intrinsically disordered C-terminal region of VIPP1 (Vc), a unique structural feature, and sought to identify its interacting protein partners. While Vc is characteristic of VIPP1 in photosynthetic organisms, recent in silico analyses of the broader VIPP1/PspA protein family have revealed that analogous regions are also present in several bacterial PspA proteins, particularly those from extremophilic species (Ma et al., manuscript under revision, included as Supplemental data). This suggests that Vc-like domains may have independently evolved multiple times as an adaptive response to environmental stressors, such as high light or oxidative conditions. Structurally, Vc is exposed on the outer surface of VIPP1 oligomers, consistent with a role in protein-protein interactions.

To identify Vc-interacting proteins, we conducted GFP-trap assays and reproducibly detected two chloroplast-localized HSP70s (cpHsc70-1 and cpHsc70-2) as well as MFP1 (Fig. 1, Fig. 2). These interactions were independently confirmed using VIPP1-His pulldown assays (Fig. 3), ruling out the possibility that the C-terminal GFP tag interfered with native protein interactions. Additionally, DnaJ-like proteins (C73 and A7) were detected in the VIPP1-His affinity purification (Supplemental Dataset 4). The presence of C73 corroborates findings from previous studies and further confirms the association between the HSP70 chaperone system and VIPP1 in Arabidopsis(48). Our results align with findings from proximity labeling studies in *Chlamydomonas* (47), where CDJ2 (a DnaJ-like protein) was identified as an interactor of VIPP1, though CGE was not. These observations suggest that full assembly of the HSP70 chaperone complex may be transient or require cofactors such as ATP. While cpHSP70 interactions were expected, our identification of MFP1 as a novel Vc interactor is particularly intriguing. MFP1 has been previously shown to interact with PTST2, a starch biosynthesis regulator, and VIPP1 has been detected in immunoprecipitates of related complexes (53, 60).

Comparison with the study by Yilmazer et al. (2023) revealed overlapping and distinct interactors. In addition to cpHSP70 and DnaJ-C73, their study reported six other potential interactors. Among these, VPL3/VIA1 (At5g03900), a PPR protein (At1g64430), and FLAP1 (At1g54520) were also detected in our MS data but were not specific to Vc, as they were present in both VIPP1-GFP and VIPP1ΔC-GFP lines(48). ALS1 (At2g31810) was detected only in *VIPP1-His/ko*, whereas a predicted transmembrane protein (At4g13220) and NADH-1 (AtCg01090) were absent or not enriched. Interestingly, lipoxygenase 2 (LOX2) was consistently detected across datasets in both *Arabidopsis* and *Chlamydomonas*, suggesting a conserved interaction with VIPP1 that does not involve Vc. The discrepancies between studies may result from differences in solubilization and immunoprecipitation conditions. Yilmazer et al. used protocols that might preserve VIPP1 oligomers differently than our methods. Our GFP-trap co-IP approach aimed to maintain oligomer integrity, which may influence which interactions are captured. Some interactions, such as with VPL3/VIA1, may involve other regions of VIPP1, such as the N-terminal domain. Moreover, the specific interaction between Vc and cpHsc70-1 may depend on VIPP1’s structural conformation—rings, rods, or filaments—which could vary between methods and conditions.

### Role of Vc in thermoprotection of TMs

Previously, we showed that deletion of Vc (*VIPP1ΔC-GFP/ko*) did not affect plant growth under normal conditions but impaired recovery of photosynthetic efficiency following heat stress, as revealed by chlorophyll fluorescence assays (21). Consistent with this, mass spectrometry profiles of *VIPP1-GFP/ko* and *VIPP1ΔC-GFP/ko* were largely similar, except for the reduced detection of cpHsc70-1 in the ΔC variant, suggesting that heat sensitivity arises from the loss of Vc–cpHsc70-1 interaction.

Supporting this, both *VIPP1ΔC-GFP/ko* and the *cphsc70-1* mutant exhibited similar heat-sensitive phenotypes in the heat-acclimation assay (49) (Fig. 4). Despite cpHsc70-1 and cpHsc70-2 shared high similarity (98% in amino acids), only *cphsc70-1* showed a strong phenotype. Because our antibody cannot distinguish between the two isoforms, whether the two proteins have functional redundancy with different accumulation levels remains unclear. Nevertheless, the fact that the *cphsc70-1/cphsc70-2* double mutant is embryo-lethal suggests their overlapping yet non-redundant functions (49). Although cpHsc70-1 protein levels were stable under heat stress, its transcript levels increased (Supplemental Fig. S3), consistent with its reported role in regulating ROS-responsive genes. cpHsc70-1 is involved in the formation of ROS by osmotic stress and regulates downstream abiotic stress-related ROS-scavenging system genes COR15A, KIN1, etc. (61). Moreover, co-silencing of *cpHsc70-1* and *cpHsc70-2* leads to *vipp1*-like phenotypes, including vesicle accumulation and chloroplast depletion. These observations imply that heat activation of cpHSP70s may occur via post-translational regulation—such as changes in stromal pH or ATP levels—rather than by increased protein synthesis. Notably, VIPP1–cpHsc70-1 interaction is enhanced by ATP depletion in vitro (45), suggesting a mechanism responsive to TM integrity.

### cpHsc70-1 governs VIPP1 disassembly and dynamics

HSP70B from *Chlamydomonas* has been implicated in modulating VIPP1 rod assembly/disassembly in vitro (44, 45), yet in vivo evidence remained elusive. The precise role of cpHSP70 in thylakostasis has never been elucidated in vascular plants. In this study, we provided conclusive evidence that cpHsc70-1 is involved in the disassembly of VIPP1 oligomers. Using VIPP1-GFP complementation lines and cpHSP70 mutants in *Arabidopsis*, we observed FVPs in both *VIPP1-GFP* and *VIPP1ΔC-GFP* lines (Fig.1A, Fig. 6). These particles likely comprise bundled filaments that remodel dynamically in response to membrane stress (41). However, *VIPP1ΔC-GFP* formed larger, static oligomers with reduced remodeling capacity, a phenotype mimicked in the *cphsc70-1* mutant (Fig. 6).

We propose a static/dynamic model in which cpHsc70-1 drives transitions between VIPP1 oligomeric states (Fig. 7). In the static state, VIPP1 forms bundled FVPs. Upon membrane stress, cpHsc70-1 associates with Vc on the filament surface, initiating disassembly (dynamic state). The liberated VIPP1 molecules then relocalize and insert into damaged membrane regions, destabilized or even raptured by stress, where they can reassemble into functional oligomers. When equilibrium is restored, VIPP1 returns to the static FVP state. Disruption of cpHsc70-2 had little effect, underscoring the specificity of cpHsc70-1 in this process. Interestingly, this model parallels the VPS4–ESCRT-III system, where ATP-driven disassembly is mediated by interactions between MIT domains (Microtubule Interacting and Trafficking domain) and MIM motifs (MIT Interacting Motifs) (62-64). Although HSP70 lacks a MIT domain, it may similarly interact with Vc to drive disassembly. Alternatively, ATP/GTPase activity of VIPP1 may contributes to its remodeling dynamics, but it warrants further investigation.

**Fig. 7.**
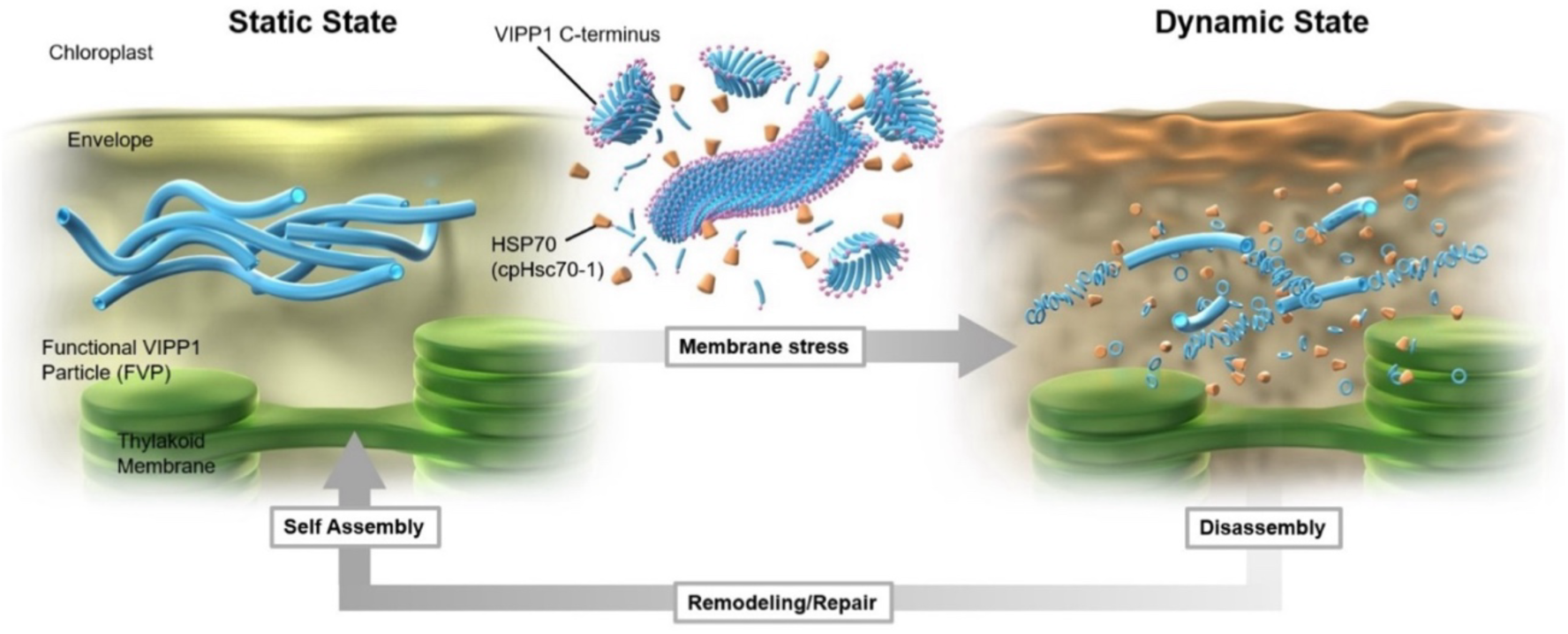
Proposed model for the role of cpHsc70-1 in VIPP1 oligomer disassembly in Arabidopsis. In the static state (left), most VIPP1 oligomers (FVPs) exist as bundled filaments, often associated with the chloroplast envelope and stroma thylakoids. Under membrane stress conditions, such as heat, VIPP1 transitions to a dynamic state (right), in which FVPs are disassembled into smaller particles by the action of cpHSP70 - predominantly cpHsc70-1 - facilitating membrane remodeling and repair on the stromal side. Based on in vitro analysis, VIPP1 can self-assemble into large FVPs of various morphologies without the need for assembly factors. However, disassembly of FVPs into smaller oligomers requires the activity of cpHSP70.

### cpHsc70-1 is required for stability of GFP-tagged proteins

VIPP1-GFP can rescue the *vipp1* phenotype, but this complementation depends on cpHsc70-1. Although cpHsc70-1 has not been structurally resolved in association with the TOC–TIC translocon or Ycf2–FtsHi import machinery (65-68), previous studies have reported its interaction with several TOC–TIC components (56, 69). In *Chlamydomonas*, loss of TIC214 impairs transit peptide processing (70), suggesting a role for cpHsc70-1 in maintaining protein import efficiency into chloroplasts. Its chaperone activity may also stabilize exogenous proteins like GFP. In *cphsc70-1* mutants, VIPP1-GFP levels were markedly reduced, particularly in true leaves(71), implying a developmental regulation of import or stabilization mechanisms. Importantly, this effect was not unique to VIPP1; CURT1A-GFP in this study and GUN1-GFP in the previous report exhibited similar instability in *cphsc70-1* mutants (59, 72). However, only VIPP1-GFP displayed impaired FVP dynamics and hypertrophy, indicating that these phenotypes are specific to VIPP1’s function.

## Conclusion

We identified Vc as a key domain mediating interaction between VIPP1 and cpHsc70-1, particularly under heat stress. This interaction facilitates ATP-dependent disassembly of VIPP1 oligomers, drawing parallels with the cytosolic ESCRT-III system. Beyond VIPP1, cpHsc70-1 also plays a broader role in stabilizing exogenous proteins, contributing to chloroplast protein quality control. Our findings highlight VIPP1 as a central player in thylakostasis, integrating protein homeostasis and membrane remodeling through its dynamic regulation by the chloroplast HSP70 system.

## Materials and Methods

### Plant materials and growth condition

*Arabidopsis thaliana* ecotype Columbia (Col) was used in this study as the wild type. Two mutants *cphsc70-1* (SALK_140810) and *cphsc70-2* (SALK_095715) were obtained from Hsou-min Li (Institute of Molecular Biology, Academia Sinica). Transgenic lines expressing VIPP1-GFP and VIPP1ΔC-GFP in *vipp1-ko,* termed as *VIPP1-GFP/ko* and *VIPP1ΔC-GFP/ko* respectively, have been described in our previous work (12). Other transgenic lines used in this study, *cphsc70-1 VIPP1ΔC-GFP/vipp1-ko* (+/+ or +/-), *cphsc70-1 VIPP1-GFP/vipp1-ko* (+/+ or +/-)*, cphsc70-1 CURT1A-GFP/Col, cphsc70-2 VIPP1ΔC-GFP/vipp1-ko* (-/-), and *cphsc70-2 VIPP1-GFP/vipp1-ko* (-/-) were created by genetic crosses, and each genotype was confirmed by subsequent genotyping (Fig. S4). The genotypes corresponding to the presence of *VIPP1-GFP* (or *VIPP1ΔC-GFP*) and *vipp1-ko* were determined as described (21). Each genotype for *cphsc70-1* and *cphsc70-2* was determined by PCR with the following primers: 1E2-S (5’-GATTCACAGAGGACAGCTAC -3’) and 1E8-AS (5’-TGAATCTCCTGATGAAGCAC -3’) for the wild-type cpHsc70-1, 70-2F (5’-CAAGCTGTTGTTAATCCGGAG-3’) and 70-2R (5’-AGCAACCATGTTTTACATGGC-3’) for the wild-type cpHsc70-2, SALK Lba1 primer (5’-TGGTTCACGTAGTGGGCCATCG-3’) and 1E8-AS (5’-TGAATCTCCTGATGAAGCAC -3’) for the presence of T-DNA insertion in *cphsc70-1*, LBb1.3 (5’-ATTTTGCCGATTTCGGAAC-3’) and 70-2R for the presence of T-DNA insertion in cpHsc70-2. The insertion of CURT1A-GFP was detected by primers Curt1A-GFP-F (5’-ATGGCGATATCAGTAGCAGCTTCGT-3’) and Curt1A-GFP-R (5’-ATGCCGTTCTTCTGCTTGTC-3’).

Seeds were sterilized by 5% (v/v) sodium hypochloride and were plated on Murashige and Skoog (MS) medium which contained 1% (w/v) sucrose. After 3 days of stratification at 4°C, MS plates were transferred to growth chamber (23°C, 50 μmol m-2 s-1 16-h photoperiod). Seeds that have cphsc70-1 mutation were plated on MS medium containing 0.5% (w/v) sucrose instead of 1%, to visualize the leaf phenotype (49). Heat shock experiment was performed essentially as described previously (49) with some modifications. Briefly, the heat shock regime was as indicated in Fig 4A, exposing 7d-old plants to 37.5 °C for 1 h, followed by 2 days for recovery and re-exposing them at 44.5 °C for 40 min. After recovery stage, seedlings were harvested and subjected to measuring chlorophyll extracted in 80% (v/v) acetone. The chlorophyll content was determined by spectrophotometry as described (73).

### Protein extraction and immunoblotting analysis

To purify chloroplasts by percoll step gradients, leaves from 30-day old plants were homogenized by blenders in isolation buffer (50 mM (w/v) HEPES/KOH, pH 7.0, 400 mM (w/v) mannitol, 1 mM (w/v) MgCl_2_, 2 mM (w/v) EDTA, 0.5% (w/v) BSA). After filtering by two layers of Miracloth (Millipore, USA), the homogenate was centrifuged at 600 *g* for 5 min at 4 °C. The pellet was resuspended in isolation buffer gently and then loaded onto 10%, 40% and 80% (v/v) Percoll step gradient (GE Healthcare, UK), followed by centrifugation at 600 *g* for 5 min at 4°C. The fractions containing chloroplasts (40%/80% interface) or broken thylakoids (10%/40% interface) were collected and washed 3 times by isolation buffer. To extract total membrane leaf proteins, seedling samples were frozen in liquid nitrogen and ground by pestles. Total leaf protein was resuspended by extraction buffer (50 mM (w/v) HEPES/KOH, pH 7.5, 330 mM (w/v) sorbitol, 5 mM (w/v) MgCl_2_, 10 mM (w/v) NaCl). After centrifugation at 15000 *g* for 10 min at 4 °C, the pellet was resuspended by extraction buffer and then denatured by SDS-sample buffer (50 mM (w/v) Tris-HCl, pH 6.8, 2% (w/v) SDS, 10% (v/v) glycerol, 0.005% (w/v) bromophenol blue, and 50 mM (w/v) dithiothreitol). The samples were subjected directly to SDS-PAGE.

The protein samples were denatured at 95 °C for 5 min and then separated by 12.5% SDS– PAGE using the conventional Laemmli (Tris-glycine) system (74) and transferred onto PVDF membranes. The resolved gels were stained with Coomassie Brilliant Blue R250 or Ponceau S as loading control. Polyclonal antibodies against VIPP1 were generated as described previously (32). Polyclonal antibodies against CURT1A was provided from Chanhong Kim (Chinese Academy of Sciences). For detecting cpHSP70, polyclonal antibodies against the pea stromal Hsp70, S78 (75), generated by Mitsuru Akita (Ehime University, Japan) was used. For GFP, a commercially available polyclonal antibody was purchased (MBL, Japan). Following immune reaction, the membranes were incubated with anti-IgG secondary antibodies (rabbit). Immunoblot signals were visualized using the Immobilon Crescendo Western HRP Substrate (Millipore, USA). The chemiluminescence signal was detected by a ChemiDoc XRS+ imaging system (Bio-Rad, USA). Protein bands on the gel were visualized by staining with Coomassie Brilliant Blue Stain One (Nacalai Tesque, Kyoto, Japan) or with a silver staining kit KANTOIII (Kanto Chemical Corp) in accordance with the manufacturers’ protocols.

### Co-immunoprecipitation analysis

To perform GFP trap analysis, purified chloroplasts from *VIPP1ΔC-GFP/vipp1-ko* and *VIPP1-GFP/vipp1-ko* (containing 3.2 mg Chl/mL) were resuspended in lysis buffer (20 mM (w/v) HEPES/KOH pH 7.0, 500 mM (w/v) NaCl, 1 mM (w/v) EDTA). After centrifugation at 6000 *g* for 10 min at 4 °C, the pellet was resuspended and solubilized by lysis buffer containing 1% (w/v) n-Dodecyl-β-D-Maltopyranoside (DDM) (w/v) by rotating for 30 min at 4 °C. The cleared lysate after centrifugation at 15000 *g* for 15 min was incubated with 25 µL of equilibrated GFP-Trap agarose beads (magnetic particles M270, chromotek, USA) by rotating for 60 min at 4 °C. The beads were sedimented by a magnet stand from the supplier and washed in lysis buffer containing 0.03% DDM for three times. The washed beads were resuspended with 2 × SDS-sample buffer to dissociate immunocomplexes, and the final eluate was analyzed by SDS-PAGE.

To perform co-immunoprecipitation using VIPP1 and cpHSP70 antibodies, each antibody was covalently coupled with Magnosphere MS160/Tosyl beads (JSR, Japan) according to the manufacturer’s protocol. Chloroplast proteins solubilized by DDM were incubated with the antibodies coupled with equilibrated magnosphere beads in binding buffer (0.1 M (w/v) borate buffer, pH9.5) by shaking at 37 °C for 18 h, followed by blocking the reaction by adding blocking reagent (10% (w/v) BSA) and further shaking at 37 °C for 6 h. The mixture was placed in the magnetic stand to discard the supernatant, followed by washing the beads for 3 times in washing buffer (25 mM (w/v) Tris-HCl, pH 7.2, 0.15 M (w/v) NaCl, 0.05% (v/v) Tween-20). The final beads were suspended in 1 × TBS-T and subjected to SDS-PAGE and western blotting.

### Transgenic lines expressing VIPP1-His and CURT1A-GFP

To create a transgenic line expressing VIPP1-His, the plasmid construction pGreen0029-VIPP1-GFP (12) was used as a backbone and modified in this study so as to generate a new construction (pGreen0229-VIPP1-His), in which the nucleotide sequence corresponding to GFP was replaced with those corresponding to His-tag (32). In pGreen0229-VIPP1-His, *VIPP1* contained truncated sequences corresponding to additional eight (EQKLISEEDL) and 6xHis peptides at the C-terminus. This construction was introduced to *Agrobacterium tumefaciens* strain GV3101, which was then infiltrated into heterozygous *vipp1-ko* (+/-) plants. In the subsequent T1 generation, we selected transformants whose genotypes were homozygous for *vipp1* (-/-) and expressing VIPP1-His. The resulting transgenic line was named *VIPP1-His/vipp1-ko* and used for protein purification using Ni-NTA agarose (GE Helthcare). The solubilized proteins extracted from chloroplasts of *VIPP1-His/vipp1-ko* were first incubated with Ni-NTA agarose at 4°C for 2 to 2.5 h. After unbound fraction was removed as flowthrough, the Ni-NTA column was washed with 25 mM imidazole-containing buffer (25 mM (v/v) imidazole, 20 mM (w/v) Tris-HCl, pH 8, 500 mM (w/v) NaCl), followed by a stepwise elution of VIPP1-His by a series of imidazole-containing buffer with different imidazole concentrations. The obtained fraction containing VIPP1-His was confirmed by SDS-PAGE and following silver staining, then subjected to MS analysis and immunoprecipitation analysis. Whenever necessary, the eluted fractions were centrifugated (7,500g at 4°C) for more than 2 h with AmiconUltra-4 (10K; Merck Millipore) to concentrate protein fraction and/or exchange buffers. The concentration of protein was estimated using the Bradford Protein Assay Kit (Bio-Rad Laboratories).

To generate transgenic plants expressing CURT1A-GFP, the plasmid pGreen0029 was digested with the restriction enzymes SacI and KpnI to insert the CURT1A-CDS and sGFP. After Agrobacterium-mediated transformation of Arabidopsis thaliana ecotype Col, transgenic plants expressing CURT1A-GFP were identified and selected.

### Mass spectrometry analysis

Eluted fractions from the GFP trap analysis were solubilized in SDS sample buffer and subsequently fractionated by 12.5% SDS-PAGE and silver staining. Elution samples from *VIPP1-GFP/vipp1-ko* and *VIPP1ΔC-GFP/vipp1-ko* were run side-by-side, and the gel areas visually detected as bands were excised from the gels. The gel bands containing proteins were dehydrated by using centrifugal concentrator (TAITEC, Japan) and then submitted for MS (MS1).

As for MS using the protein fraction purified from *VIPP1-His/vipp1-ko*, the purified fraction was directly used. To select proteins specifically found in *VIPP1-His/vipp1-ko*, the same protein fraction purified from the wild-type Col through Ni-NTA column was also subjected to MS (MS2), and the proteins only detected in *VIPP1-His/vipp1-ko* was included in the dataset. Gel slices from silver-stained gels were de-stained, reduced, alkylated and tryptically digested according to established protocols (76). Afterwards, peptides were desalted using in-house made C18 STAGE-tips as described (77), dried by vacuum centrifugation and stored at -80°C until further use. Samples were analysed on an LC-MS/MS system consisting of an Ultimate 3000 nanoLC (Thermo Fisher Scientific) coupled to a Q Exactive Plus mass spectrometer (Thermo Fisher Scientific) via a Nanospray Flex ion source (Thermo Fisher Scientific). Peptide samples were first dissolved in a solution of 2% acetonitrile and 0.05% trifluoroacetic acid (LB) and loaded onto a trap column (Acclaim PepMap 100 C18, dimensions: 300 µm × 5 mm, particle size: 5 µm, pore size: 100 Å) using LB at 10 µl/min for 3 minutes. The peptides were then separated on a reversed-phase column (PepSep 15, 75 µm × 15 cm, particle size: 1.9 µm, pore size: 100 Å, Bruker) at 250 nl/min. The eluents for separation were 0.1% formic acid in ultrapure water (A) and 80% acetonitrile with 0.1% formic acid in ultrapure water (B). The gradient was programmed as follows: 5% to 8% B over 2 minutes, 8% to 45% B over 33 minutes, 45% to 99% B over 2 minutes, and 99% B for 5 minutes.

Survey scans (MS1) covered m/z 350-1400 in positive ion mode with 70,000 resolution (at m/z 200). The AGC target was 3e6 with a maximum injection time of 50 ms. For MS2 analysis, the 12 most abundant ions (charge states 2-4) underwent HCD fragmentation at 27% normalized collision energy. MS2 parameters were: AGC target of 5e4, maximum injection time of 80 ms, intensity threshold 1e4, and a precursor isolation window of 1.5 m/z. Dynamic exclusion was set to 15 s.

The protein and peptide identification and label-free quantification were performed using MaxQuant (version 2.0.3.0) (78). Raw MS files were searched against the Arabidopsis TAIR10 reference proteome with default MaxQuant settings. The search incorporated variable modifications (methionine oxidation and protein N-terminal acetylation) and a fixed modification (cysteine carbamidomethylation). A 1% false discovery rate threshold was applied for both peptides and proteins.

### Confocal laser scanning microscopy

For observation of chloroplasts in living leaf tissues, a piece of Arabidopsis leaf was excised and examined using a Confocal laser scanning microscopy (LAS-X, Leica Stellaris5). Three fluorescence channels were utilized: chlorophyll, GFP, and Rhodamine B. For chlorophyll autofluorescence, excitation was achieved at 488 nm, with emission detected at 650–700 nm. GFP fluorescence was excited at 488 nm, with emission detected at 500–550 nm. Rhodamine B was excited at 543 nm, with emission collected at 570–620 nm. The envelope membrane staining is performed by infiltrating the leaves with Rhodamine B solution (10 µM (w/v) Rhodamine B, 150mM sorbitol, 20mM HEPES/KOH, pH8.0) 5 times and then letting them stand for 2 mins

To quantify VIPP1-GFP particles (FVP: functional VIPP1 particles), the number of FVP per chloroplast and the longest length of each particle was measured by ImageJ (Fiji) software (version 1.51n). Data was collected from at least 3 independent images. GFP particles numbers and length were measured manually with each mutant containing around 100 individuals. Time lapse videos were recorded at 8 frames/s for more than 20 seconds.

## Author Contributions

W.S. conceived the research and designed the experiments with D.L.; D.L. performed most of the experiments under the supervision of W.S., including co-immunoprecipitation, heat-shock experiments, generation and characterization of mutants related to *cphsc70-1* and *cphsc70-2*, and microscopic observations; S.W.G. created and characterized *VIPP1-His/vipp1-ko*; S.O., M.S. and M.H. performed mass spectrometric analysis along with W.S. and D.L.; D.L. and W.S. analyzed data; all authors confirmed the experimental data; D.L. and W.S. wrote the manuscript with inputs from the coauthors.

## Classification

Plant Biology

## Other Supplemental Data

Supplemental Datasets 1 to 4

Supplemental Videos 1 t

## Acknowledgments

We thank Tsuneaki Takami, Alexandre Muhire, Fiona Wacera Wahinya, and Rie Hijiya for their technical assistance in the Sakamoto laboratory. We also thank Mitsuru Akita (Ehime University) for providing HSP70 antibodies, Chanhong Kim (Chinese Academy of Sciences) for providing CURT1A antibodies, Hsou-min Li (Academia Sinica) for *cphsc70-1* and *cphsc70-2* seeds. This work was supported by the KAKENHI Grant from the Ministry of Education, Culture, Sports, Science and Technology (23H04959 to W.S. and S.O.) and from the Japan Society for the Promotion of Science (21H02508 and 24K02044 to W.S.). M.H. acknowledges funding by the Deutsche Forschungsgemeinschaft (DFG HI739/13-3), and the RECTOR program in association with Okayama University. D.L. is supported by the scholarship from Ohara Foundation.

## Supplemental data captions

Supplemental Dataset 1. List of proteins detected by mass spectrometry in Fig. 1*C*.

Supplemental Dataset 2. List of proteins detected by mass spectrometry in Suppl Fig. S1.

Supplemental Dataset 3. List of proteins detected by mass spectrometry in Suppl Fig. S2.

Supplemental Dataset 4. List of proteins detected by mass spectrometry in Fig. 3.

Supplemental Video 1. A video showing VIPP1 dynamics in true leaves from *VIPP1-GFP/ko*.

Supplemental Video 2. A video showing VIPP1 dynamics in true leaves from *VIPP1ΔC-GFP/ko*.

Supplemental Video 3. A video showing VIPP1 dynamics in true leaves from *hsc70-2 VIPP1-GFP/ko*.

Supplemental Video 4. A video showing VIPP1 dynamics in true leaves from *hsc70-2 VIPP1ΔC-GFP/ko*.

Supplemental Video 5. A video showing VIPP1 dynamics in cotyledons from *VIPP1-GFP/ko*.

Supplemental Video 6. A video showing VIPP1 dynamics in cotyledons from *VIPP1ΔC-GFP/ko*.

Supplemental Video 7. A video showing VIPP1 dynamics in cotyledons from *hsc70-1 VIPP1-GFP/Col*.

Supplemental Video 8. A video showing VIPP1 dynamics in cotyledons from *hsc70-1 VIPP1ΔC-GFP/ko*.

---

**Figure S1.**
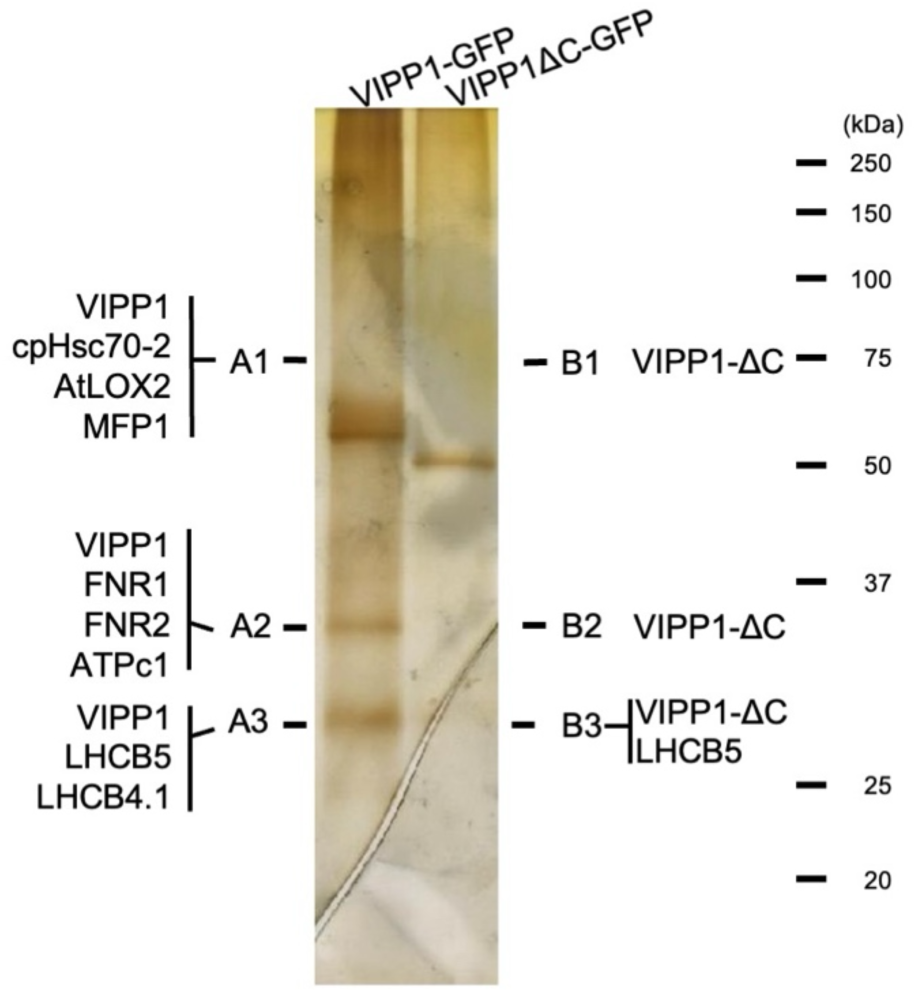
Mass spectrometry analysis of immunoprecipitated proteins with VIPP1-GFP and VIPP1ΔC-GFP (Replicate 2). Solublized chloroplast protein extracts from *VIPP1-GFP* and *VIPP1ΔC-GFP* were immunoprecipitated with anti-GFP serum-coupled beads. The precipitated proteins were separated by SOS-PAGE and visualized by silver staining. Gel slices in *VIPP1-GFP* (A1-A3) and *VIPP1ΔC-GFP* (B1-B3) were excised and analyzed by mass spectrometry. Identified proteins are presented in Supplemental Dataset 2.

**Figure S2.**
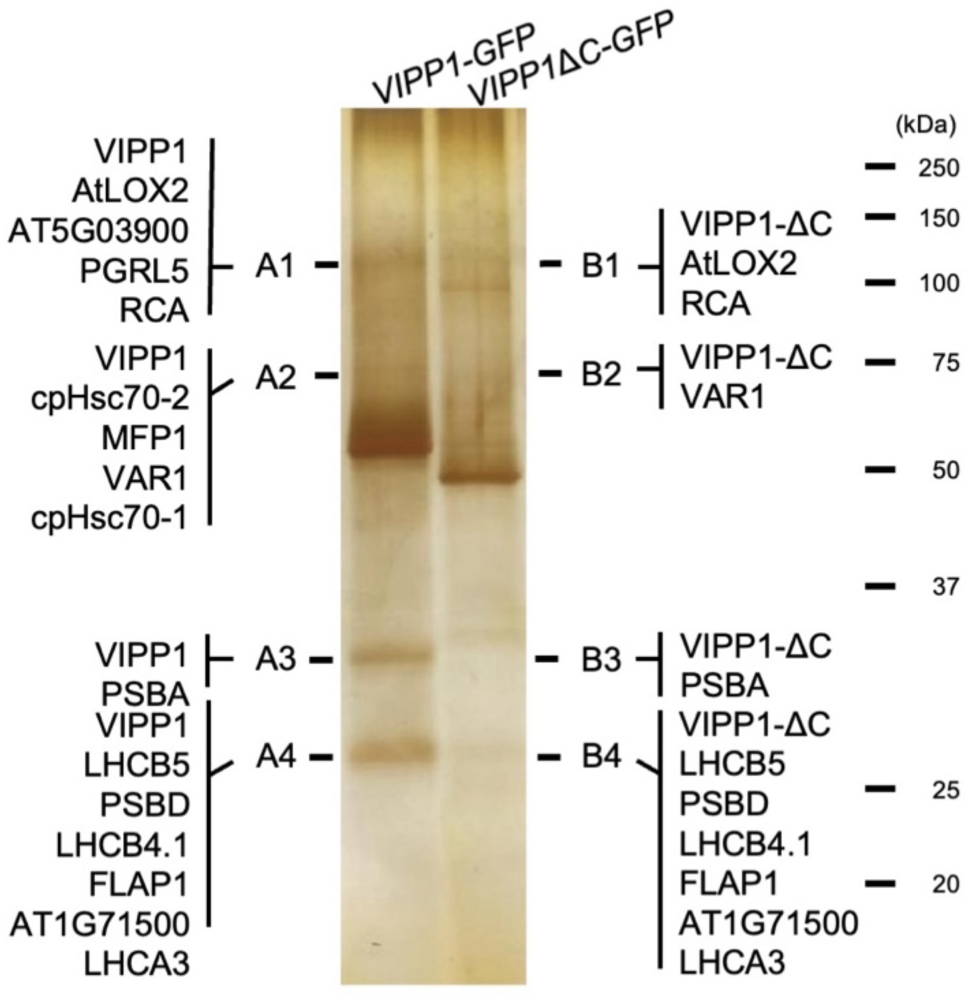
Mass spectrometry analysis of immunoprecipitated proteins with VIPP1-GFP and VIPP1ΔC-GFP (Replicate 3). Solublized chloroplast protein extracts from *VIPP1-GFP* and *VIPP1ΔC-GFP* were immunoprecipitated with anti-GFP serum-coupled beads. The precipitated proteins were separated by SDS-PAGE and visualized by silver staining. Gel slices in *VIPP1-GFP* (A1-A4) and *VIPP1ΔC-GFP* (B1-B4) were excised and analyzed by mass spectrometry. Identified proteins are presented in Supplemental Dataset 3.

**Figure S3.**
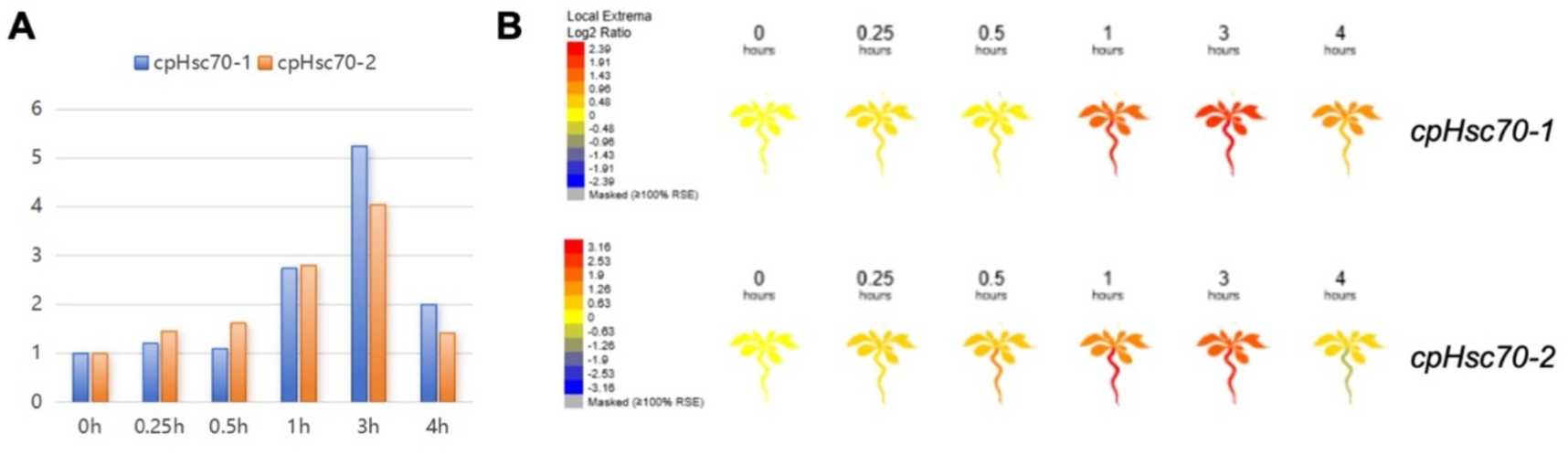
Induction of *cpHsc70-1* and *cpHsc70-2* RNA levels after heat shock. Data were obtained from the ePlant database (https://bar.utoronto.ca/eplanU). (A) Transcript levels of *cpHsc70-1* and *cpHsc70-2* follwing heat shock at 38 °C. (B) Schematic representation of the heat-induced expression profiles of *cpHsc70-1* and *cpHscl0-2*.

**Figure S4.**
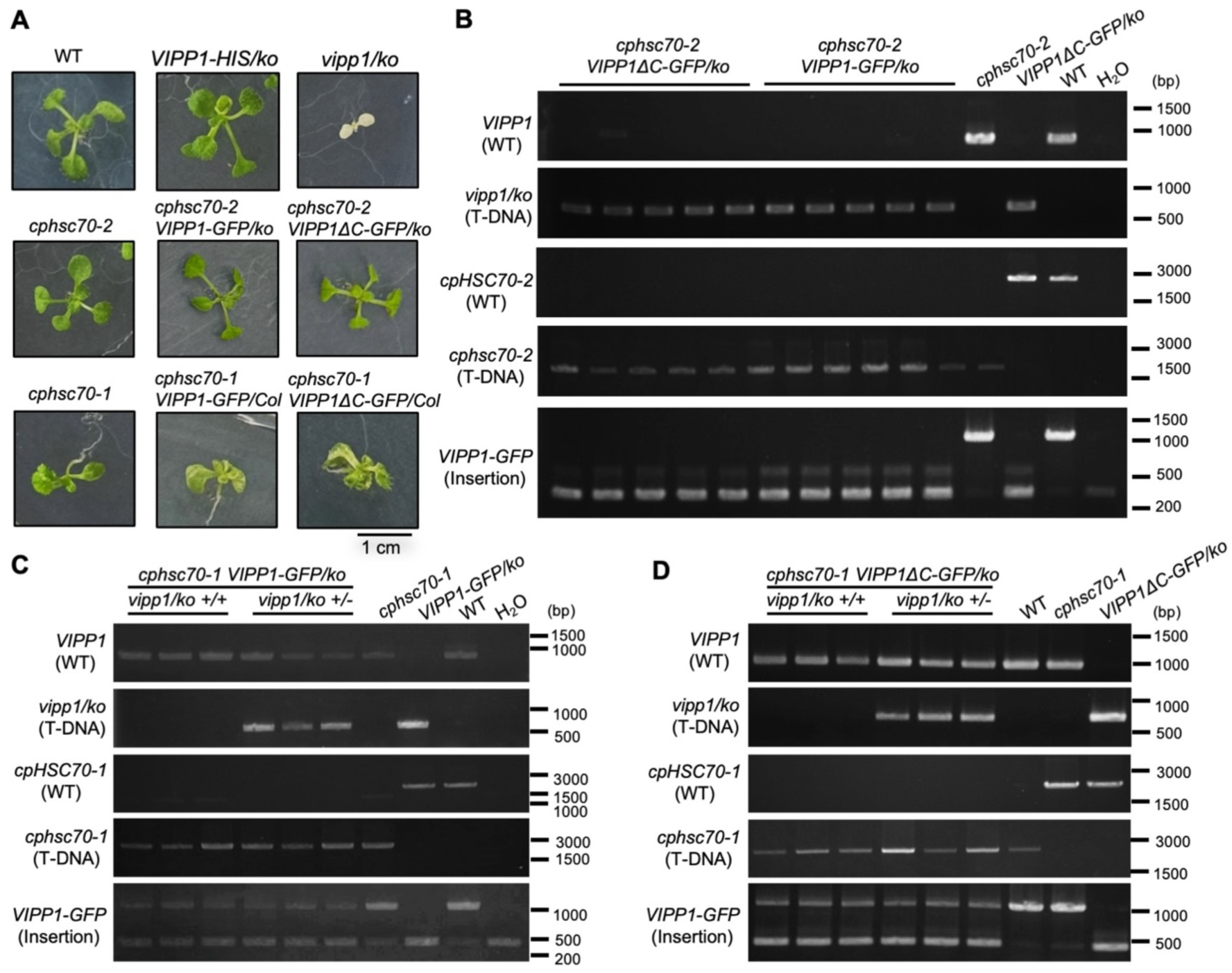
PCR-based genotyping of mutant lines used in this study. (A) Seedling phenotype of the indicated mutants. Genotypes are labeled above each image. (B) Verification of the genotypes corresponding to *VIPP1-GFP* insertion, *vipp1-ko,* and cphsc70-2 alleles (indicated on the left) in the mutant lines indicated at the top. Three independent lines for *cphscl0-2 VIPP1ΔC-GFP/ko* and *cphscl0-2 VIPP1-GFP/ko* are shown. (C) Verification of the genotypes corresponding to *VIPP1-GFP* insertion, *vipp1-ko,* and cphsc70-1 alleles (indicated on the left) in the mutant lines indicated at the top. Three independent lines homozygous for *vipp1-ko* +/+ and heterozygous for *vipp1-ko* +/-are shown. (D) As in (C), but using the transgenic line *VIPP1ΔC-GFP* instead of *VIPP1-GFP.* The relative sizes of DNA fragments in each agarose gel are shown on the right, based on molecular markers.

**Figure S5.**
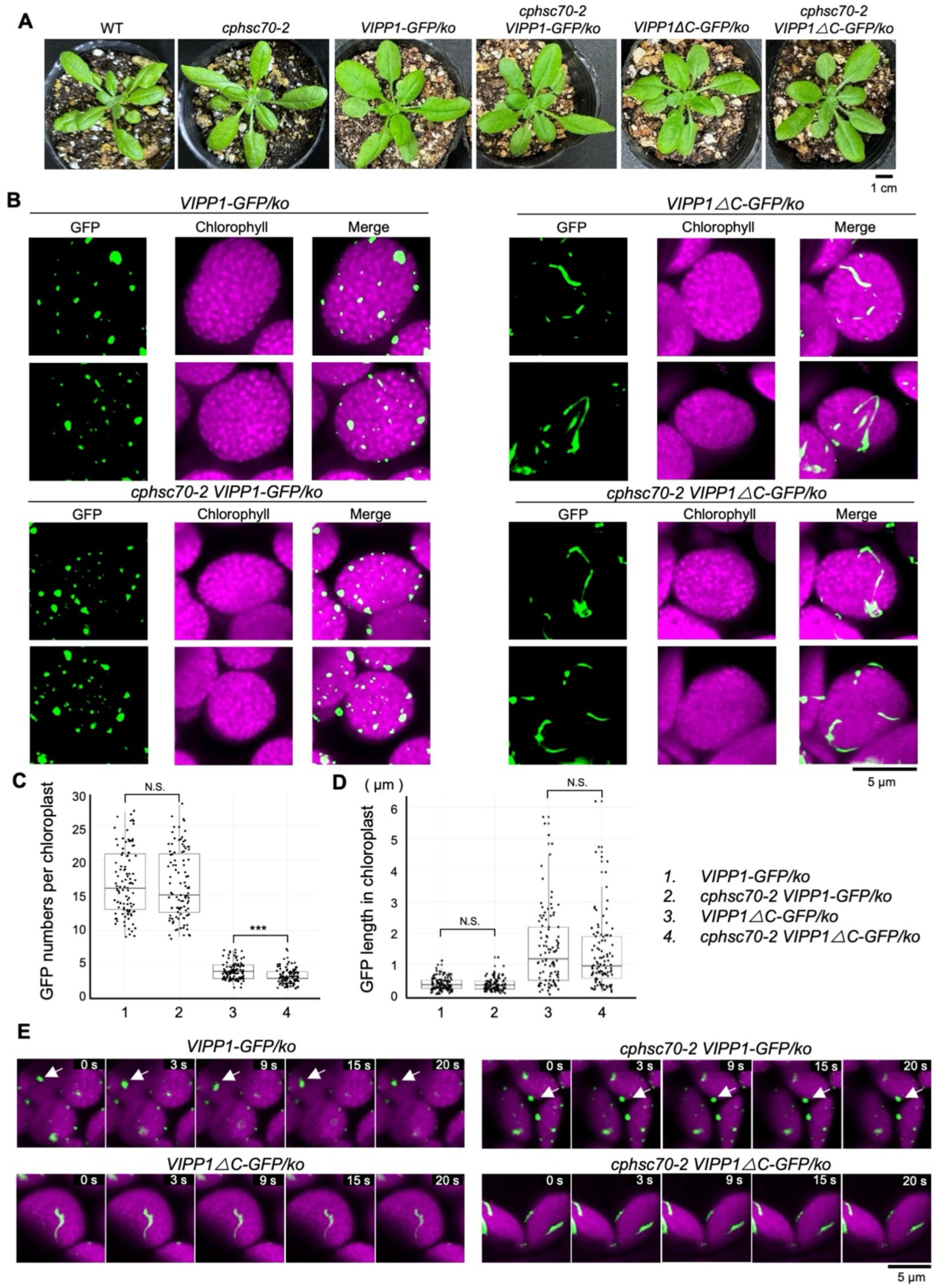
Confocal microscopic observation of GFP-tagged VIPP1 in chloroplasts. (A) Plant morphologies at 30 days after germination. Genotype are indicated above the images. (B) Representative images of GFP-tagged VIPP1 in true leaf chloroplasts. Top left; VIPP1-GFP in *vipp1-ko,* Bottom left; VIPP1-GFP in *vipp1-kplcphsc70-2,* Top right; VIPP1ΔC-GFP in *vipp1-ko,* Bottom right; VIPP1ΔC-GFP in *vipp1-kp/cphsc70-2.* GFP fluorescence, chlorophyll autofluorescence, and merged images are shown. Bar=S µm. (C) Box plots showing the number of GFP particles per chloroplast for the indicated genotypes (N > 104). (D) Box plots showing the length (longest axis) of individual GFP particles, measured using image J (N > 106). (E) Time lapse images of GFP-tagged VIPP1 particles in true leaves from *VIPP1-GFP/ko* (top left), *VIPP1ΔC-GFP/ko* (bottom left), *cphsc70-2 VIPP1-GFP/ko* (top right), and *cphsc70-2 VIPP1ΔC-GFP/ko* {bottom right). Shown are five time points (0-20 seconds) from each time-lapse series. White arrows indicate fast moving GFP particles.

## References

1. S. Eberhard, G. Finazzi, F. A. Wollman, The dynamics of photosynthesis. Annu. Rev. Genet. 42, 463–515 (2008).

2. Z. Adam, D. Charuvi, O. Tsabari, R. R. Knopf, Z. Reich, Biogenesis of thylakoid networks in angiosperms: knowns and unknowns. Plant. Mol. Biol. 76, 221–234 (2011).

3. M. Rantala, S. Rantala, E. M. Aro, Composition, phosphorylation and dynamic organization of photosynthetic protein complexes in plant thylakoid membrane. Photochem. Photobiol. Sci. 19, 604–619 (2020).

4. M. Pribil, M. Labs, D. Leister, Structure and dynamics of thylakoids in land plants. J. Exp. Bot. 65, 1955–1972 (2014).

5. J. M. Anderson, P. Horton, E. H. Kim, W. S. Chow, Towards elucidation of dynamic structural changes of plant thylakoid architecture. Philos. Trans. R. Soc. Lond. B Biol. Sci. 367, 3515–3524 (2012).

6. W. S. Chow, E. H. Kim, P. Horton, J. M. Anderson, Granal stacking of thylakoid membranes in higher plant chloroplasts: the physicochemical forces at work and the functional consequences that ensue. Photochem. Photobiol. Sci. 4, 1081–1090 (2005).

7. A. J. Dorne, J. Joyard, R. Douce, Do thylakoids really contain phosphatidylcholine? Proc. Natl. Acad. Sci. U.S.A. 87, 71–74 (1990).

8. I. Sakurai et al., Lipids in oxygen-evolving photosystem II complexes of cyanobacteria and higher plants. J. Biochem. 140, 201–209 (2006).

9. U. Armbruster et al., Arabidopsis CURVATURE THYLAKOID1 proteins modify thylakoid architecture by inducing membrane curvature. Plant Cell 25, 2661–2678 (2013).

10. M. Patil, S. Seifert, F. Seiler, J. Soll, S. Schwenkert, FZL is primarily localized to the inner chloroplast membrane however influences thylakoid maintenance. Plant Mol. Biol. 97, 421–433 (2018).

11. Y. Ogawa, M. Iwano, T. Shikanai, W. Sakamoto, FZL, a dynamin-like protein localized to curved grana edges, is required for efficient photosynthetic electron transfer in Arabidopsis. Front. Plant Sci. 14, 1279699 (2023).

12. L. Zhang, Y. Kato, S. Otters, U. C. Vothknecht, W. Sakamoto, Essential role of VIPP1 in chloroplast envelope maintenance in Arabidopsis. Plant Cell 24, 3695–3707 (2012).

13. D. Kroll et al., VIPP1, a nuclear gene of Arabidopsis thaliana essential for thylakoid membrane formation. Proc. Natl. Acad. Sci. U.S.A. 98, 4238–4242 (2001).

14. P. Albanese, S. Tamara, G. Saracco, R. A. Scheltema, C. Pagliano, How paired PSII-LHCII supercomplexes mediate the stacking of plant thylakoid membranes unveiled by structural mass-spectrometry. Nat. Commun. 11, 1361 (2020).

15. A. Trotta et al., Defining the heterogeneous composition of Arabidopsis thylakoid membrane. Plant J. 121, e17259 (2025).

16. M. Ostermeier, A. Garibay-Hernández, V. J. C. Holzer, M. Schroda, J. Nickelsen, Structure, biogenesis, and evolution of thylakoid membranes. Plant Cell 36, 4014–4035 (2024).

17. R. Kobayashi, T. Suzuki, M. Yoshida, Escherichia coli phage-shock protein A (PspA) binds to membrane phospholipids and repairs proton leakage of the damaged membranes. Mol. Microbiol. 66, 100–109 (2007).

18. E. Aseeva et al., Complex formation of Vipp1 depends on its alpha-helical PspA-like domain. J. Biol. Chem. 279, 35535–35541 (2004).

19. G. Jovanovic et al., The N-terminal amphipathic helices determine regulatory and effector functions of phage shock protein A (PspA) in Escherichia coli. J. Mol. Biol. 426, 1498–1511 (2014).

20. S. Otters et al., The first α-helical domain of the vesicle-inducing protein in plastids 1 promotes oligomerization and lipid binding. Planta 237, 529–540 (2013).

21. L. Zhang, H. Kondo, H. Kamikubo, M. Kataoka, W. Sakamoto, VIPP1 Has a Disordered C-Terminal Tail Necessary for Protecting Photosynthetic Membranes against Stress. Plant Physiol. 171, 1983–1995 (2016).

22. H. M. Li, Y. Kaneko, K. Keegstra, Molecular cloning of a chloroplastic protein associated with both the envelope and thylakoid membranes. Plant Mol. Biol. 25, 619–632 (1994).

23. U. C. Vothknecht, S. Otters, R. Hennig, D. Schneider, Vipp1: a very important protein in plastids?! J. Exp. Bot. 63, 1699–1712 (2012).

24. S. Westphal, L. Heins, J. Soll, U. C. Vothknecht, Vipp1 deletion mutant of Synechocystis: a connection between bacterial phage shock and thylakoid biogenesis? Proc. Natl. Acad. Sci. U.S.A. 98, 4243–4248 (2001).

25. S. Zhang, G. Shen, Z. Li, J. H. Golbeck, D. A. Bryant, Vipp1 is essential for the biogenesis of Photosystem I but not thylakoid membranes in Synechococcus sp. PCC 7002. J. Biol. Chem. 289, 15904–15914 (2014).

26. A. Nordhues et al., Evidence for a role of VIPP1 in the structural organization of the photosynthetic apparatus in Chlamydomonas. Plant Cell 24, 637–659 (2012).

27. S. M. Lo, S. M. Theg, Role of vesicle-inducing protein in plastids 1 in cpTat transport at the thylakoid. Plant J. 71, 656–668 (2012).

28. R. Hennig et al., IM30 triggers membrane fusion in cyanobacteria and chloroplasts. Nat. Commun. 6, 7018 (2015).

29. C. McDonald, G. Jovanovic, O. Ces, M. Buck, Membrane Stored Curvature Elastic Stress Modulates Recruitment of Maintenance Proteins PspA and Vipp1. mBio 6, e01188–01115 (2015).

30. J. Theis et al., VIPP1 rods engulf membranes containing phosphatidylinositol phosphates. Sci. Rep. 9, 8725 (2019).

31. N. Ohnishi et al., Distinctive in vitro ATP Hydrolysis Activity of AtVIPP1, a Chloroplastic ESCRT-III Superfamily Protein in Arabidopsis. Front. Plant Sci. 13, 949578 (2022).

32. N. Ohnishi, L. Zhang, W. Sakamoto, VIPP1 Involved in Chloroplast Membrane Integrity Has GTPase Activity in Vitro. Plant Physiol. 177, 328–338 (2018).

33. B. Junglas, D. Schneider, What is Vipp1 good for? Mol. Microbiol. 108, 1–5 (2018).

34. L. Zhang, W. Sakamoto, Possible function of VIPP1 in maintaining chloroplast membranes. Biochim. Biophys. Acta 1847, 831–837 (2015).

35. T. K. Gupta et al., Structural basis for VIPP1 oligomerization and maintenance of thylakoid membrane integrity. Cell 184, 3643–3659.e3623 (2021).

36. J. Liu et al., Bacterial Vipp1 and PspA are members of the ancient ESCRT-III membrane-remodeling superfamily. Cell 184, 3660–3673.e3618 (2021).

37. B. Junglas et al., PspA adopts an ESCRT-III-like fold and remodels bacterial membranes. Cell 184, 3674–3688.e3618 (2021).

38. S. Pan et al., The cyanobacterial protein VIPP1 forms ESCRT-III-like structures on lipid bilayers. Nat. Struct. Mol. Biol. 32, 543–554 (2025).

39. B. Junglas et al., Structural basis for Vipp1 membrane binding: from loose coats and carpets to ring and rod assemblies. Nat. Struct. Mol. Biol. 32, 555–570. (2025).

40. S. Naskar et al., Mechanism for Vipp1 spiral formation, ring biogenesis, and membrane repair. Nat. Struct. Mol. Biol. 32, 571–584. (2025).

41. S. W. Gachie et al., The thylakoid membrane remodeling protein VIPP1 forms bundled oligomers in tobacco chloroplasts. Plant Physiol. 198, kiaf137 (2025).

42. P. H. Su, H. Y. Lin, Y. H. Lai, Two Arabidopsis Chloroplast GrpE Homologues Exhibit Distinct Biological Activities and Can Form Homo- and Hetero-Oligomers. Front. Plant Sci. 10, 1719 (2019).

43. L. A. de Luna-Valdez et al., Functional analysis of the Chloroplast GrpE (CGE) proteins from Arabidopsis thaliana. Plant Physiol. Biochem. 139, 293–306 (2019).

44. C. Liu et al., J-domain protein CDJ2 and HSP70B are a plastidic chaperone pair that interacts with vesicle-inducing protein in plastids 1. Mol. Biol. Cell 16, 1165–1177 (2005).

45. C. Liu et al., The chloroplast HSP70B-CDJ2-CGE1 chaperones catalyse assembly and disassembly of VIPP1 oligomers in Chlamydomonas. Plant J. 50, 265–277 (2007).

46. F. Gao, W. Wang, W. Zhang, C. Liu, α-Helical Domains Affecting the Oligomerization of Vipp1 and Its Interaction with Hsp70/DnaK in Chlamydomonas. Biochemistry 54, 4877–4889 (2015).

47. E. Kreis et al., TurboID reveals the proxiomes of Chlamydomonas proteins involved in thylakoid biogenesis and stress response. Plant Physiol. 193, 1772–1796 (2023).

48. I. Yilmazer, et al., A conserved ESCRT-II-like protein participates in the biogenesis and maintenance of thylakoid membranes. bioRxiv [Preprint] (2023), 10.1101/2023.10.10.561251 Accessed 10 May 2025.

49. P. H. Su, H. M. Li, Arabidopsis stromal 70-kD heat shock proteins are essential for plant development and important for thermotolerance of germinating seeds. Plant Physiol. 146, 1231–1241 (2008).

50. B. L. Lin et al., Genomic analysis of the Hsp70 superfamily in Arabidopsis thaliana. Cell Stress Chaperones 6, 201–208 (2001).

51. D. Y. Sung, E. Vierling, C. L. Guy, Comprehensive expression profile analysis of the Arabidopsis Hsp70 gene family. Plant Physiol. 126, 789–800 (2001).

52. P. A. Harder, R. A. Silverstein, I. Meier, Conservation of matrix attachment region-binding filament-like protein 1 among higher plants. Plant Physiol. 122, 225–234 (2000).

53. M. Sharma et al., MFP1 defines the subchloroplast location of starch granule initiation. Proc. Natl. Acad. Sci. U.S.A. 121, e2309666121 (2024).

54. M. S. Bae, E. J. Cho, E. Y. Choi, O. K. Park, Analysis of the Arabidopsis nuclear proteome and its response to cold stress. Plant J. 36, 652–663 (2003).

55. L. Rizhsky et al., When defense pathways collide. The response of Arabidopsis to a combination of drought and heat stress. Plant Physiol. 134, 1683–1696 (2004).

56. P. H. Su, H. M. Li, Stromal Hsp70 is important for protein translocation into pea and Arabidopsis chloroplasts. Plant Cell 22, 1516–1531 (2010).

57. V. R. Agashe, F. U. Hartl, Roles of molecular chaperones in cytoplasmic protein folding. Semin. Cell Dev. Biol. 11, 15–25 (2000).

58. J. Larkindale, E. Vierling, Core genome responses involved in acclimation to high temperature. Plant Physiol. 146, 748–761 (2008).

59. G. Z. Wu et al., Control of retrograde signalling by protein import and cytosolic folding stress. Nat. Plants 5, 525–538 (2019).

60. D. Seung, T. B. Schreier, L. Bürgy, S. Eicke, S. C. Zeeman, Two Plastidial Coiled-Coil Proteins Are Essential for Normal Starch Granule Initiation in Arabidopsis. Plant Cell 30, 1523–1542 (2018).

61. F. Ding, F. Li, B. Zhang, A plastid-targeted heat shock cognate 70-kDa protein confers osmotic stress tolerance by enhancing ROS scavenging capability. Front. Plant Sci. 13, 1012145 (2022).

62. J. McCullough, A. Frost, W. I. Sundquist, Structures, Functions, and Dynamics of ESCRT-III/Vps4 Membrane Remodeling and Fission Complexes. Annu. Rev. Cell Dev. Biol. 34, 85–109 (2018).

63. B. E. Mierzwa et al., Dynamic subunit turnover in ESCRT-III assemblies is regulated by Vps4 to mediate membrane remodelling during cytokinesis. Nat. Cell Biol. 19, 787–798 (2017).

64. E. Isono, ESCRT Is a Great Sealer: Non-Endosomal Function of the ESCRT Machinery in Membrane Repair and Autophagy. Plant Cell Physiol. 62, 766–774 (2021).

65. K. Liang et al., Structural insights into the chloroplast protein import in land plants. Cell 187, 5651–5664.e5618 (2024).

66. K. Liang et al., Conservation and specialization of the Ycf2-FtsHi chloroplast protein import motor in green algae. Cell 187, 5638–5650.e5618 (2024).

67. Z. Jin et al., Structure of a TOC-TIC supercomplex spanning two chloroplast envelope membranes. Cell 185, 4788–4800.e4713 (2022).

68. H. Liu, A. Li, J. D. Rochaix, Z. Liu, Architecture of chloroplast TOC-TIC translocon supercomplex. Nature 615, 349–357 (2023).

69. R. M. Ratnayake, H. Inoue, H. Nonami, M. Akita, Alternative processing of Arabidopsis Hsp70 precursors during protein import into chloroplasts. Biosci. Biotechnol. Biochem. 72, 2926–2935 (2008).

70. S. Ramundo et al., Coexpressed subunits of dual genetic origin define a conserved supercomplex mediating essential protein import into chloroplasts. Proc. Natl. Acad. Sci. U.S.A. 117, 32739–32749 (2020).

71. H. Tsukaya, T. Tsuge, H. Uchimiya, The cotyledon: A superior system for studies of leaf development. Planta 195, 309–312 (1994).

72. L. Tadini et al., GUN1 influences the accumulation of NEP-dependent transcripts and chloroplast protein import in Arabidopsis cotyledons upon perturbation of chloroplast protein homeostasis. Plant J. 101, 1198–1220 (2020).

73. R. J. Porra, W.A. Thompson, P.E. Kriedemann, Determination of accurate extinction coefficients and simultaneous equations for assaying chlorophylls a and b extracted with four different solvents: verification of the concentration of chlorophyll standards by atomic absorption spectroscopy. Biochim. Biophys. Acta 975, 384–394 (1989).

74. U. K. Laemmli, Cleavage of structural proteins during the assembly of the head of bacteriophage T4. Nature 227, 680–685 (1970).

75. M. Akita, E. Nielsen, K. Keegstra, Identification of protein transport complexes in the chloroplastic envelope membranes via chemical cross-linking. J. Cell Biol. 136, 983–994 (1997).

76. A. Shevchenko, H. Tomas, J. Havlis, J. V. Olsen, M. Mann, In-gel digestion for mass spectrometric characterization of proteins and proteomes. Nat. Protoc. 1, 2856–2860 (2006).

77. J. Rappsilber, M. Mann, Y. Ishihama, Protocol for micro-purification, enrichment, pre-fractionation and storage of peptides for proteomics using StageTips. Nat. Protoc. 2, 1896–1906 (2007).

78. J. Cox, M. Mann, MaxQuant enables high peptide identification rates, individualized p.p.b.-range mass accuracies and proteome-wide protein quantification. Nat. Biotechnol. 26, 1367–1372 (2008).

